# Single cell decomposition of multicellular aging programs associated with impaired lung regeneration

**DOI:** 10.1101/2025.07.24.666371

**Authors:** Janine Gote-Schniering, M. Camila Melo-Narváez, Gowtham Boosarpu, David Lauer, Sai Rama Sridatta Prakki, Huijuan Wang, Matthias Brunner, Simrah Khan, Meshal Ansari, Michael Ammeter, Eshita Jain, Nikola Puda, Konstantin Wiedemann, Carina Steinchen, Sara Ashgapour, Ariane Eceiza, Michael Dudek, Christoph H. Mayr, Yuexin Chen, Ilias Angelidis, Önder Yildirim, Percy Knolle, Gerald Burgstaller, Melanie Königshoff, Safwen Kadri, Martin Mück-Häusl, Fabian J. Theis, Mareike Lehmann, Herbert B. Schiller

## Abstract

Aging impairs the regenerative capacity of mammalian organs and is a major risk factor for organ fibrosis. Mechanisms underlying persistent fibrosis after lung injury in old individuals remain unclear. We used longitudinal single-cell RNA-seq after lung injury and dissected aging effects computationally and experimentally at baseline and during repair. In old mice, sustained fibroblast activation in the resolution phase of fibrosis was associated with prolonged epithelial senescence and persistent epithelial-mesenchymal crosstalk. Single-cell interpretable tensor decomposition analysis revealed that aging most strongly affected T/B-lymphocytes and macrophages. Notably, we identified a Granzyme K-high CD8^+^ T cell state that was unique to aged mice, co-localized with epithelial progenitors, and its co-culture or Gzmk treatments in lung organoids impaired progenitor function by inducing stem cell senescence. In summary, our study highlights the effects of immune aging on epithelial progenitor function and provides a time-resolved high resolution map of lung regeneration in the context of aging.

## Introduction

Chronic lung diseases (CLD) predominantly affect individuals over 60 years of age, feature impaired regeneration, and are the third leading cause of death worldwide^1^. The lungs are particularly vulnerable to aging effects due to their continuous exposure to environmental stressors. Given the rising incidence of CLD such as chronic obstructive pulmonary disease (COPD) and pulmonary fibrosis (PF) with age, it is key to understand how aging affects lung regeneration. Tissues from CLDs exhibit multiple hallmarks of aging, including stem cell exhaustion, telomere attrition, and cellular senescence, suggesting that (premature) aging and impaired regenerative capacity are a key driver of CLD disease onset and progression.

Functionally, the aged organism shows a reduced capacity to recover from stress, including lung injury. Even in the absence of injury, aging induces substantial changes in the cellular and molecular composition of lung tissues^2,3^ with the local immune system undergoing particularly profound remodeling^4^. Immune aging, including thymic involution and clonal restriction of lymphocytes, alters pulmonary surveillance, with resident memory CD8⁺ T cells accumulating in dysfunctional states that sustain low-grade inflammation rather than resolution^5–9^. Recent observations also suggest that hematopoietic shifts, including bone marrow-derived myeloid biases, perpetuate fibrotic cascades by delaying macrophage maturation and IL-10 signaling^10^.

Regenerative processes in the lung rely on transient injury-associated regenerative cell states (RCSs), such as Krt8+ alveolar differentiation intermediates (Krt8-ADI) and Cthrc1⁺ myofibroblasts, which bridge injury response and restoration but can persist maladaptively in diseased tissues^11–15^. Experimental models like bleomycin-induced injury in aged mice at least partially mimic disease relevant processes in human PF. While young mice typically resolve bleomycin-induced experimental fibrosis, aged mice exhibit persistent fibrotic remodeling associated with decreased myofibroblast apoptosis and increased oxidative stress also seen in patients^16^. Generally, aged animals seem to display delayed resolution and persistent RCSs, at least partially driven by metabolic shifts and unresolved hypoxia^14^.

It remains poorly characterized which aspects of lung aging and functional cellular interactions control the emergence, function, and resolution of these RCSs. Advanced computational tools, such as single cell tensor decomposition, now enable dissection of coordinated programs across cell types, revealing age-specific covariation of cell states across individuals^17^. To address better the time resolved aspects of cellular dynamics and leverage new opportunities for systems-level analysis, we conducted longitudinal single-cell RNA sequencing (scRNA-seq) on young and aged mouse lungs at baseline and post-bleomycin (days 0–37), spanning the early inflammatory, fibrogenic, and resolution phases of tissue repair. Employing interpretable tensor decomposition we unraveled aging-linked multicellular programs, including prolonged senescence and epithelial-mesenchymal signaling that sustained fibroblast activation in the resolution phase of aged animals. In particular, we identified aging specific states of T and B lymphocytes and macrophages, exemplified by a unique Gzmk-high CD8⁺ T cell state, and its effect on progenitor function and the persistence of senescence-like pro-fibrotic cell states.

## Results

### Longitudinal scRNA-seq reveals age-dependent changes in cellular dynamics upon lung injury

To elucidate the cellular and molecular mechanisms underlying aging-associated impairments in lung regeneration, we performed longitudinal scRNA-seq on whole-lung suspensions from 55 young (3 months) and old (18 months) mice across six time points following bleomycin injury (0, 3, 10, 20, 30, and 37 days post-injury; dpi) (**Fig. 1A**). The selected time points captured key stages of the injury-repair process, including acute inflammation (3 dpi), fibrogenesis (10 and 20 dpi), and the resolution phase of fibrosis (30 and 37 dpi) (**Fig. 1B**).

**Figure 1.**
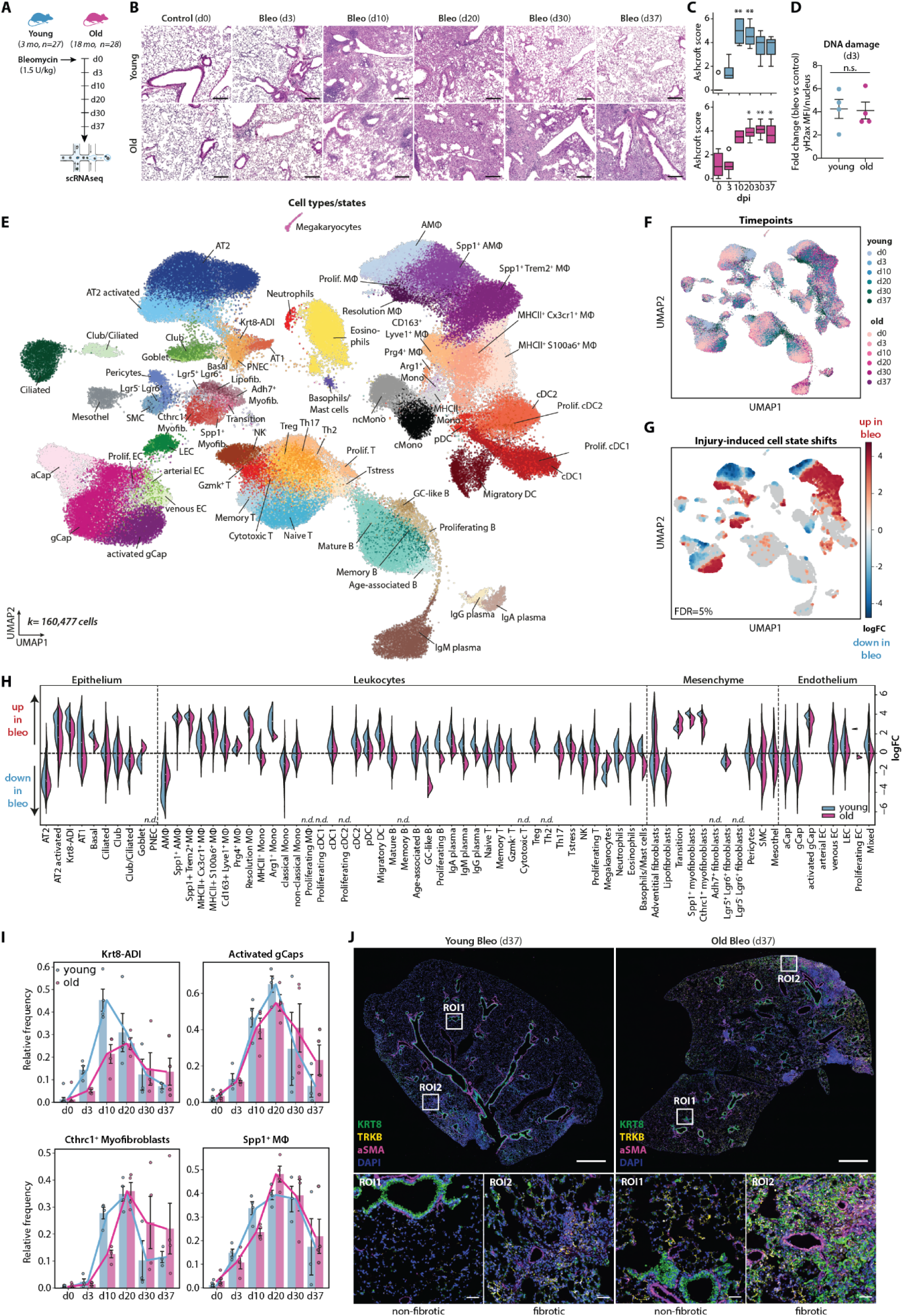
A longitudinal scRNA-seq atlas reveals age-dependent changes in cellular regeneration dynamics following bleomycin-induced lung injury. (A) Schematic overview of the experimental design. (B) Representative H&E-stained lung tissue images from young and aged mice at the investigated post-injury time points (0–37 dpi). Scale bars = 100 µm. (C) Time-resolved semiquantitative analysis of the fibrosis extent using Ashcroft scores in young (upper panel) and aged (lower panel) mice. Statistical analysis: Kruskal–Wallis test with Dunn’s post hoc test. (D) Relative induction of γH2AX mean fluorescence intensity per nucleus in young and aged mice compared to controls at 3 dpi. (E, F) UMAP embedding of the complete single-cell dataset (160,477 cells) annotated by cell type/state (69 identified states, E) or by time point (F). (G, H) Milo analysis identifying differential cellular neighborhoods in bleomycin-treated versus control mice. (G) UMAP representation highlighting neighborhoods upregulated (red) or downregulated (blue) in bleomycin-treated mice. (H) Beeswarm plot of differentially abundant neighborhoods mapped to cell type annotations, showing log-fold changes in young versus aged mice following bleomycin injury. n.d. = not determined/neighborhood mapping to mixed cluster. (I) Time-resolved changes in the relative abundance of multicellular regenerative cell states (RCSs) in young and aged mice. Mean trajectories and individual sample bar plots with standard error of the mean (SEM) are shown. (J) Representative 4i image of lung tissue from young and aged mice at 37 dpi showing persistent Krt8-ADI cells, TRKB⁺ activated gCap cells, and αSMA⁺ myofibroblasts in fibrotic and non-fibrotic areas of aged lungs (scale bar for overview image = 1 mm; ROI1/2, scale bars = 50 µm). n = 3 biological replicates.

Histopathological analysis revealed that young mice developed a robust yet transient fibrotic response, with Ashcroft scores peaking at 10 dpi and progressively resolving until 37 dpi. In contrast, old mice exhibited a delayed fibrotic peak around 20 dpi, with fibrosis scores remaining elevated until 37 dpi (**Fig. 1C**). These findings suggest that aging alters the regenerative trajectory, leading to sustained fibrotic remodeling rather than efficient repair. To exclude that these differences in regenerative capacity and fibrosis progression were due to variations in initial injury severity, we quantified bleomycin-induced DNA damage by staining for γH2AX foci at 3 dpi (**Fig. 1D**). The levels of DNA damage were comparable between young and aged mice, confirming that the observed differences between young and old mice at later timepoints were not driven by disparities in initial injury severity but rather by aging-associated alterations in lung repair mechanisms (**Fig. 1D**).

We generated a longitudinal single cell atlas of age-dependent lung regeneration by profiling 160,477 high-quality single-cell transcriptomes spanning all major lung cell lineages and conditions (**Fig. 1E/F, Supplementary Fig. 1A-E, Datafile S1,** https://hschillerlabshiny.shinyapps.io/BleomycinAging/). Hierarchical annotation defined 18 meta-cell types (**Supplementary Fig. 1B**) corresponding to 69 distinct cell types and states, each characterized by unique marker gene signatures (**Fig. Supplementary Figure 1F / Data file S2**).

Bleomycin injury led to profound compositional shifts in cell types and states across epithelial, stromal, endothelial, and immune compartments in both young and aged mice, as assessed by differential cell abundance testing using *Milo*^18^ (**Fig. 1G/H)** and confirmed by relative frequency analysis **(Supplementary Fig. 1G).** These shifts included the induction of recently described lung injury-associated regenerative cell states (RCSs), such as *Krt8*⁺ alveolar differentiation intermediate (Krt8-ADI) cells, *Cthrc1*⁺ myofibroblasts, and *Spp1*⁺ alveolar and interstitial macrophages (**Fig. 1H)**. Additionally, we identified an injury-activated general capillary (gCAP) population marked by *Ntrk2, Sparcl*, and *Vwa1* expression, which was recently implicated as induced capillary state (iCAP) in dysregulated vascular repair following viral injury and aging^14,19^ (**Fig. 1H**).

Given that RCSs transiently emerge during lung regeneration, we next examined their time-resolved dynamics in both young and aged mice (Fig. 1I, Supplementary Fig. 2). Compartment-specific subcluster analysis revealed distinct temporal patterns in RCSs expansion (Fig. 1I). While multilineage RCSs were induced in both groups, their early-stage expansion (10 dpi) was significantly attenuated in aged mice. Furthermore, in contrast to the transient nature of these populations in young lungs, aged mice exhibited persistence of a subset of RCSs, including Krt8-ADI (Supplementary Fig. 3), *Ntrk2*⁺ activated gCaps (Supplementary Fig. S4) and *Cthrc1^+^*myofibroblasts (Supplementary Fig. 5). Of note, this age dependent difference was not observed for *Spp1*⁺ macrophages, which resolved similarly in both groups (Supplementary Fig. 6). To better understand the impact of this prolonged RCSs presence in aged mice, we examined their spatial localization using multiplexed immunofluorescence (4i, Fig. 1J, Supplementary Fig. 2A/B). While in young mice at 37 dpi, epithelial, stromal and endothelial RCSs were predominantly confined to fibrotic areas, old mice exhibited persistent RCSs also in, minimally injured areas, suggesting a maladaptive regenerative program where failure to resolve these RCSs contributed to sustained fibrosis in aged lungs.

### Aging Alters Molecular Programs Underlying Injury Resolution

To investigate the molecular basis of age-dependent alterations in cellular regeneration dynamics, we performed differential gene expression analysis by diffxpy (Wald test) and quantified the magnitude of the aging effects per cell type using *Augur*^20^. Specifically, we analyzed transcriptional differences at the meta-cell type level between young and aged mice at baseline (0 dpi), as well as time-resolved, injury-induced expression changes by comparing transcriptomes at 0 dpi with corresponding post-injury time points for each age group (Supplementary Fig. 7A**/B**, **Datafile S3-5**). Even in the absence of injury, aged lungs exhibited marked transcriptional changes across multiple cellular compartments compared to young lungs, with alveolar epithelial cells showing the highest degree of age-associated alterations in the *Augur* analysis (Supplementary Fig. 7A). Upon injury, both young and aged mice displayed dynamic gene expression changes across several cellular compartments, with the most pronounced quantitative responses observed in cells of the alveolar cell circuit, including macrophages, fibroblasts, alveolar epithelial cells, and capillary endothelial cells (CECs) (Supplementary Fig. 7B).

To gain deeper insight into the molecular programs underlying divergent regenerative outcomes in young versus aged mice, we examined the temporal expression patterns of genes differentially expressed between young and aged lungs at 37 dpi. Using unsupervised clustering of pseudobulk expression profiles, we identified 9 gene clusters showing time-dependent activation and 11 clusters showing suppression in aged lungs, respectively (Fig. 2A**/B, Supplementary** Fig. 7C**/D**). Gene Ontology (GO) and MSigDB hallmark pathway enrichment analysis of these gene clusters revealed distinct pathway profiles associated with aging-related impaired injury resolution (Fig. 2C**/D, Supplementary** Fig. 7E**/F**). This included sustained inflammatory responses, such as type I and II interferon signaling (cluster 1). These pathways were primarily activated in immune cells such as dendritic cells (DCs) and monocytes, as confirmed by pathway signature scoring at the meta-cell type level (Fig. 2D**, Supplementary** Fig. 7F). Notably, excessive and prolonged interferon signaling has been shown to disrupt lung repair following influenza infection^21^ and its age-associated upregulation may contribute to the inflammatory environment in aged lungs^22^.

**Figure 2.**
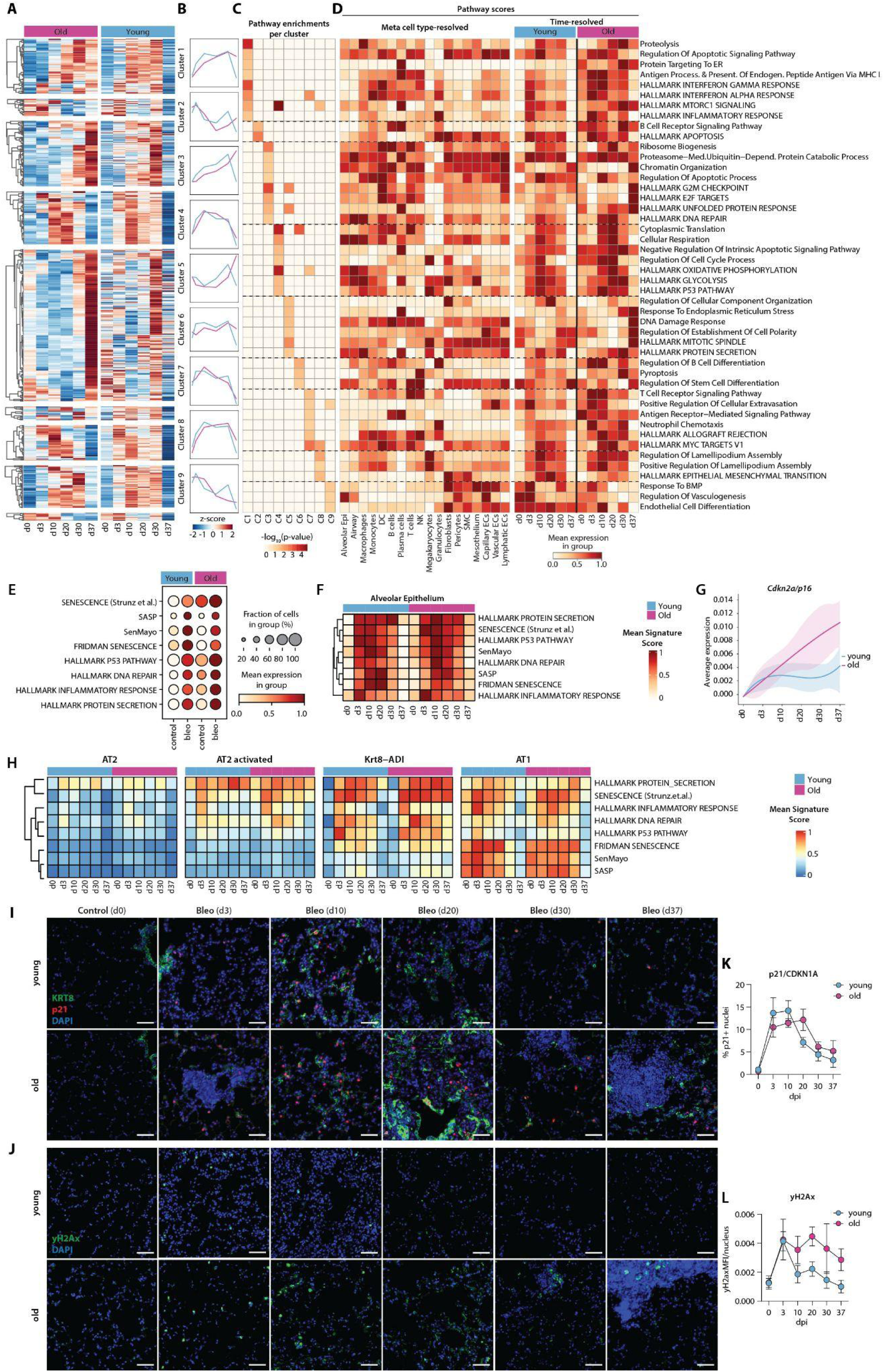
Aging alters molecular programs underlying injury resolution. (A, B) Unsupervised clustering of temporal gene expression profiles based on the 847 upregulated DEGs between young and aged mice at 37 dpi, analyzed at the pseudobulk level. (A) Heatmap showing gene expression profiles across the nine identified clusters. (B) Average gene expression trajectories of the nine clusters over time. (C) GO enrichment analysis of cluster-specific gene profiles. (D) Time-resolved gene signature scoring of the identified pathways at the meta cell type level. Scaled average expression is shown. (E) Senescence-associated gene signature scoring across the entire dataset, stratified by treatment and age. (F, H) Time- and age-resolved scoring of senescence-associated gene signatures in the alveolar epithelium overall (F), and separately by individual alveolar epithelial cell states (H). (G) Smoothed average expression of *Cdkn2a* (*p16*) over time in young and aged mice. (I, J) Representative immunofluorescence images of lung tissue from young (top panels) and aged mice (bottom panels) at 0, 3, 10, 20, 30, and 37 dpi, stained for KRT8 (green) and CDKN1A/p21 (red) (I) or γH2AX (green) (J). Scale bar = 50 µm. (K, L) Quantification of the percentage of p21⁺ nuclei (K) and γH2AX mean fluorescence intensity per nucleus (L) in young versus aged mice. *n* = 3 biological replicates. Error bars indicate the SEM.

In addition, we identified sustained activation of pathways associated with molecular aging hallmarks, particularly those linked to proteostasis dysfunction, such as the unfolded protein response, ER stress, and proteasome-mediated ubiquitin catabolic processes (clusters 3 and 5). Dysregulated epithelial proteostasis has been previously implicated in pulmonary fibrosis pathobiology, as it disrupts alveolar regeneration by blocking AT2-to-AT1 differentiation and driving the persistence of Krt8-ADI^23–25^.

Furthermore, we observed sustained, cell type-specific activation of multiple senescence-associated pathways, including DNA damage and repair signaling, cell cycle regulation (e.g., G2/M checkpoint, E2F target genes), and protein secretion programs. Senescence signature scoring based on curated, publicly available gene sets revealed upregulation of cellular senescence signatures in aged compared to young mice, both at baseline and following injury (Fig. 2E, Supplementary Fig. 8A). We and others have previously demonstrated that senescence of alveolar epithelial cells and fibroblasts is a key driver of impaired lung regeneration and fibrogenesis^26–31^. In line with these findings, senescence-associated pathways remained persistently activated within the alveolar epithelial and fibroblast compartments in aged lungs (Fig. 2F**, Supplementary** Fig. 8B-D), with the strongest enrichment observed in Krt8-ADI cells and myofibroblast subpopulations (Fig. 2H, Supplementary Fig. 8E). This was further supported by sustained elevation of *Cdkn2a*/*p16* alveolar mRNA expression following injury in aged lungs (Fig. 2G). Immunofluorescent staining for DNA damage and the canonical senescence marker p21/CDKN1A confirmed persistent senescence and DNA damage responses in aged lungs post-injury. (Fig. 2I**-L**)).

These findings suggest that age-associated dysregulation of cellular stress responses, chronic inflammation, and persistent senescence are associated with failure of regenerative resolution and the progression of fibrosis in aged lungs.

### Persistence of Krt8-ADI-Myofibroblast crosstalk in aged lungs

We next sought to determine how these divergent gene programs during aging translate into differential cell-cell communication patterns. We modelled the injury-induced cellular interaction networks using NicheNet, an approach that infers functional interactions based on prior knowledge about ligand-receptor interactions and their downstream transcriptional consequences. We observed a higher number of ligand-receptor interactions in young compared to aged mice throughout the entire time course, suggesting that aging is associated with a global reduction in intercellular communication (Supplementary Fig. 9A**/B**). Throughout the entire injury-repair time course, we identified persistent interactions between macrophages, fibroblasts, and alveolar epithelial cells, forming a key signaling axis for fibrogenesis. Notably, in young mice, these interactions dynamically resolved over time, while in aged mice, these interactions were persistently dysregulated, characterized by fibroblast-inward signaling that failed to resolve, reflecting the non-resolving fibrosis at the tissue level (Fig. 3A**, Supplementary** Fig. 9A).

**Figure 3.**
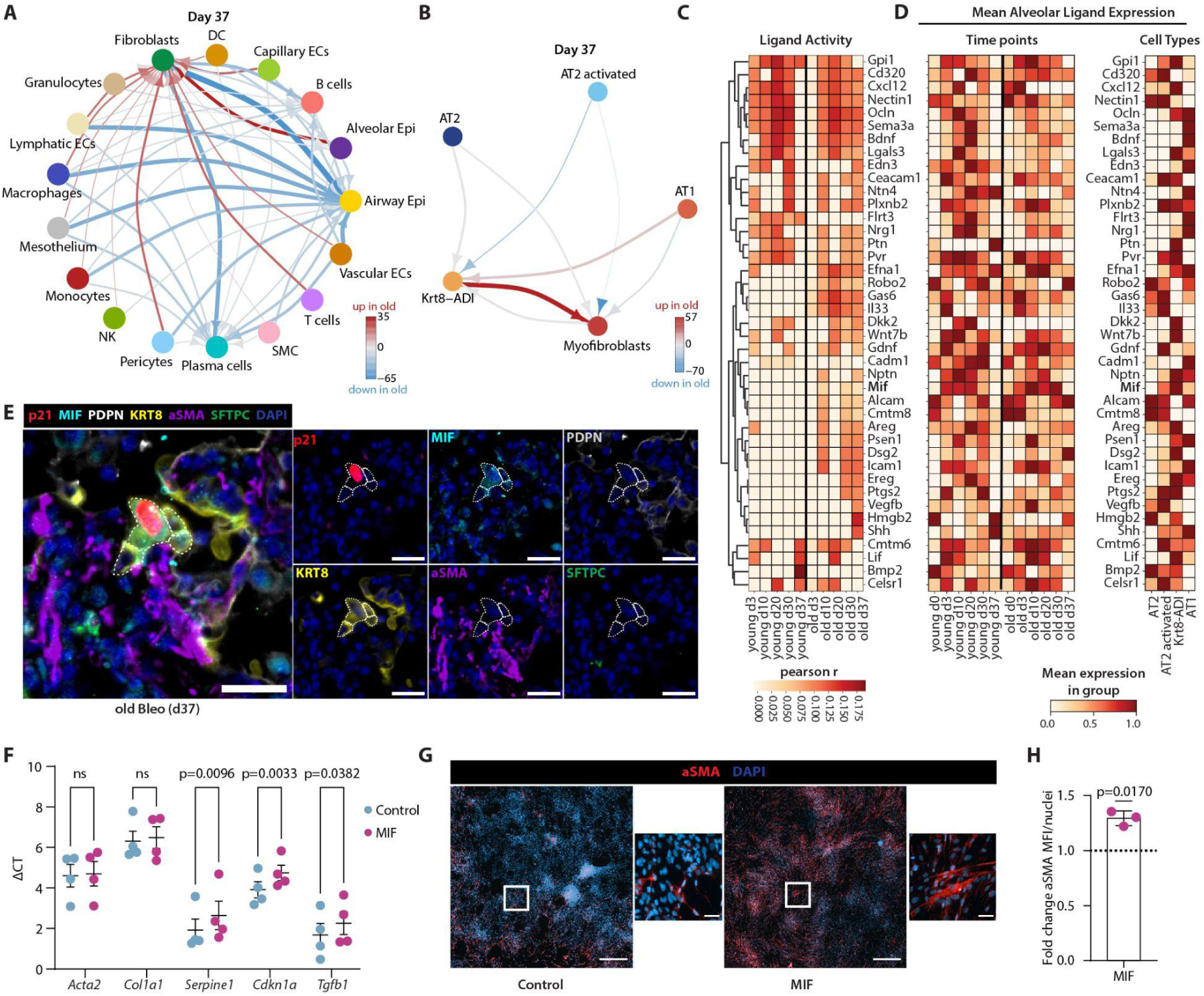
Aging is linked to persisting Krt8-ADI-Myofibroblast crosstalk. (A, B) NicheNet-based analysis of injury-induced cell–cell communication at day 37 post-injury comparing young and aged mice. Increased signaling in aged mice is shown in red; decreased interactions are shown in blue. (A) Differential signaling interactions at the meta cell type level. (B) Targeted analysis of interactions between alveolar epithelial cell states and myofibroblasts. (C) Heatmap of predicted ligand activity scores for alveolar cell-derived ligands uniquely predicted to activate myofibroblasts at day 37 in aged mice. Pearson correlation coefficients (r) are shown. (D) Heatmaps showing the average expression of the predicted myofibroblast-activating ligands in the entire alveolar compartment over time (left) and resolved by alveolar cell state (right). Scaled average expression is shown. (E) Representative 4i image of lung tissue from aged mice at day 37 post-injury, showing accumulation of MIF⁺/p21⁺/KRT8⁺ ADI cells in close proximity to αSMA⁺ myofibroblasts. Scale bars = 25 µm. n = 3 biological replicates. (F) qPCR analysis of fibrotic marker gene expression in MIF-treated versus mock-treated primary lung fibroblasts. ΔCT values are shown; error bars represent SEM. Statistical analysis: two-way ANOVA with Sidak’s multiple comparisons test. n = 4 biological replicates. (G) Representative immunofluorescence images showing αSMA expression in MIF- or vehicle-treated primary fibroblasts. Scale bar = 500 µm; zoom-in = 50 µm. (H) Quantification of relative induction of αSMA mean fluorescence intensity per nucleus in MIF-versus vehicle-stimulated fibroblasts. n=3 biological replicates. Statistical analysis: Paired Wilcoxon test.

Recent studies have demonstrated that a coordinated and transient alveolar-mesenchymal crosstalk between Krt8-ADI and myofibroblasts is critical for successful lung regeneration, while its persistence can promote fibrogenesis^13^. We next investigated whether dysregulation of this alveolar-mesenchymal crosstalk underlies age-associated failure of injury resolution and applied targeted NicheNet modeling of functional interactions between alveolar epithelial cell subtypes and myofibroblasts (Fig. 3B**, Supplementary** Fig. 9B). Indeed, our analysis suggested that persistent Krt8-ADI-to-myofibroblast crosstalk might drive sustained fibroblast activation in aged mice at 37 dpi (Fig. 3B). Focusing on the specific ligands predicted to mediate this sustained Krt8-ADI–myofibroblast interaction at 37 dpi in aged animals, we identified 40 unique ligands, several of which have previously been implicated in lung fibrosis (Supplementary Fig. 9C). Examination of the trajectory of ligand activity over time (Fig. 3C) revealed that 13 ligands were exclusively predicted throughout the entire disease course in aged mice, but not in young mice, including EREG and PTGS2, two ligands highly expressed in Krt8-ADI cells that have been previously described to drive myofibroblast activation and fibrosis progression^32,33^. In addition to these, we identified Macrophage Migration Inhibitory Factor (MIF) as a potential driver of myofibroblast activation specifically in aged mice across the entire time course (Fig. 3C). MIF was increased in aged alveolar epithelial cells compared to young counterparts at all time points, with Krt8-ADI cells identified as the main cellular source of MIF within the alveolar epithelium (Fig. 3D). Given that MIF is a known component of the senescence-associated secretory phenotype (SASP)^34,35^, its elevated expression in aged mice ties into our earlier findings that Krt8-ADI cells in aged lungs exhibit sustained senescence phenotypes. To confirm the predicted NicheNet cellular crosstalk *in situ*, we performed multiplexed immunofluorescence (4i) and found Krt8-ADI cells (KRT8⁺, proSPC⁻) co-expressing the canonical senescence marker CDKN1A/p21 and high levels of MIF directly adjacent to αSMA⁺ myofibroblasts in aged mice in fibrogenic zones at 37 dpi (Fig. 3E).

To further functionally validate the impact of MIF on fibroblast activation, we stimulated lung fibroblasts with recombinant MIF in vitro. This stimulation resulted in increased expression of profibrotic genes, including *Col1a1*, *Serpine1*/PAI-1, and *Tgfb1* (Fig. 3F), as well as increased αSMA protein expression, a key marker of myofibroblast differentiation in primary human lung fibroblasts (Fig. 3G**/H**).

To further determine whether MIF also affects lung regenerative potential, we treated lung organoid cultures with recombinant MIF. This treatment led to a significant reduction in both colony-forming efficiency and organoid size, indicating that MIF not only activates fibroblasts but also negatively impacts alveolar epithelial regeneration (Supplementary Fig. 9D-F).

These results suggest that MIF is sufficient to drive fibroblast activation and may reduce stem cell potential, thereby contributing to sustained fibrosis in aged lungs.

### Aging leads to reshaping of the myeloid and lymphoid cell compartments

To analyze how aging affects multicellular gene programs that may predispose to failed regenerative outcomes we used single-cell Independent Tensor Decomposition (*scITD*), that identifies common axes of interindividual variation by considering joint expression variation across multiple cell types^17^ (Fig. 4A). We identified five distinct latent factors that showed significant associations with age, bleomycin treatment, and injury time points (Fig. 4A, Supplementary Fig. 10A-D). One of these factors (factor 1) corresponded to injury-specific multicellular responses, which was enriched in mice at the peak of fibrosis, particularly at 10 and 20 dpi, and was driven by coordinated gene expression changes in alveolar epithelial cells, fibroblasts, capillary ECs, and macrophages and remained activated in aged mice through 37 dpi (**Fig 4B-C, Supplementary** Fig. 10A). In addition to this injury-specific factor, we identified a distinct aging-associated factor (factor 4) that was significantly enriched in aged mice at all time points. This factor was primarily driven by gene expression changes in T and B lymphocytes and macrophages, suggesting that immune dysregulation plays a central role in age-associated regenerative failure (Fig. 4A**-D**).

**Figure 4.**
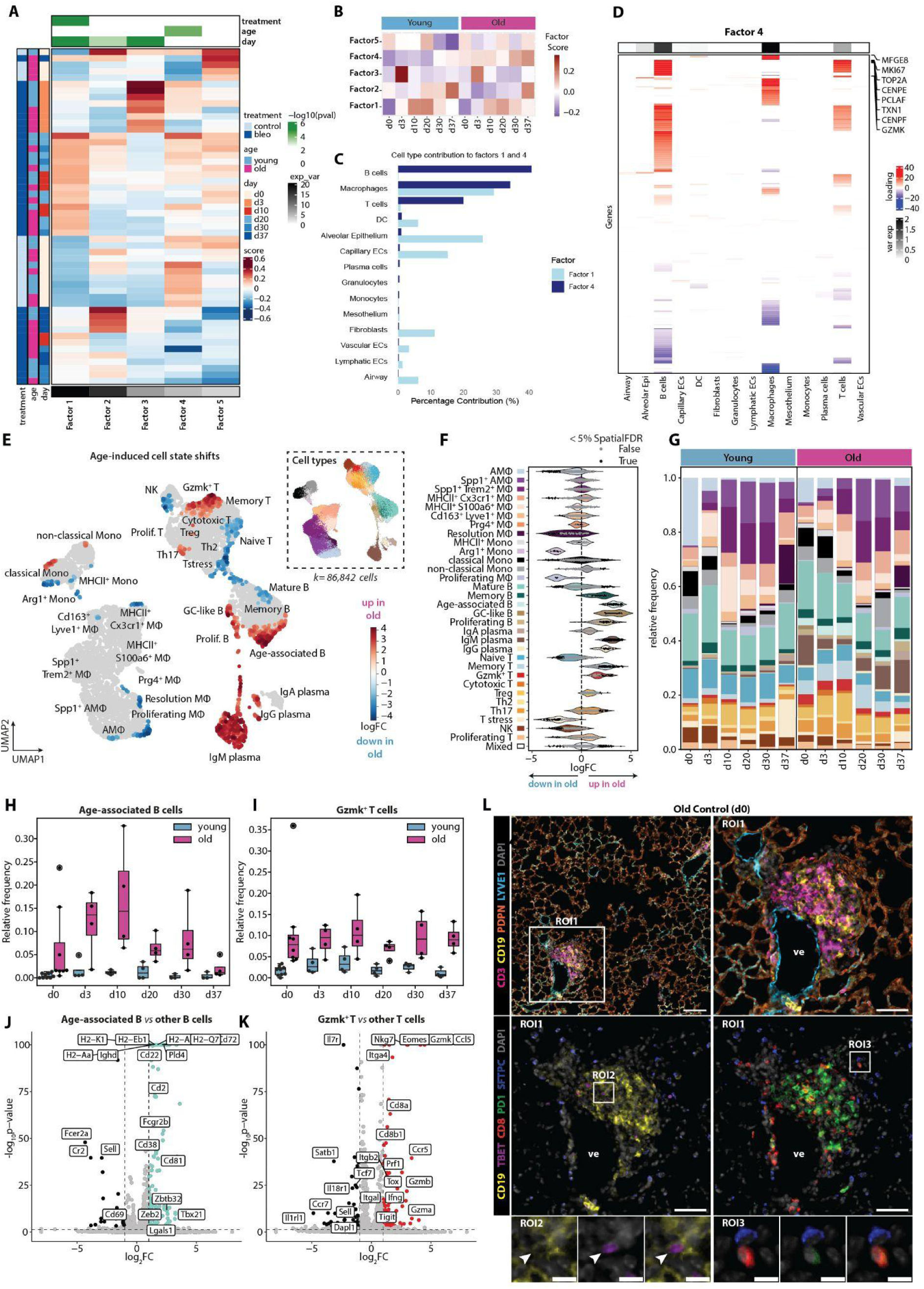
Lung aging is associated with immune compartment remodeling and the emergence of age-specific lymphocyte populations. (A) Heatmap summary of scITD analysis. (B) Time- and age-resolved heatmap of scITD factor scores (top panel) and corresponding cell type contributions (bottom panel). (C) Cell types contributions (percentage) to the factors 1 and 4 (D) Loading matrix for scITD factor 4, identifying key gene contributors. (E,F) Milo analysis showing differential cellular neighborhoods in the macrophage/monocyte and T/B lymphocyte compartment between young and aged lungs. (E) UMAP representation highlighting neighborhoods upregulated in aged mice (red) and downregulated (blue) relative to young mice. (F) Beeswarm plot mapping differentially abundant neighborhoods to cell type annotations, showing log-fold changes in young versus aged mice. (G) Time-resolved compositional analysis of immune cell types in lungs from young and aged mice. (H, I) Time-resolved changes in the relative abundance of age-associated B cells (H) and Gzmk⁺ T cells (I). (J, K) Volcano plots showing differentially expressed genes (DEGs) between age-associated B cells and other B cells (J), and between Gzmk⁺ T cells and other T cells (K). (L) Representative 4i image of lung tissue from aged mice showing accumulation of CD19⁺/TBET⁺ age-associated B cells and CD8⁺/PD1⁺ Gzmk⁺ T cells within tertiary lymphoid-like structures composed of CD3⁺ T cells and CD19⁺ B cells, and adjacent to Lyve1⁺ blood vessels (scale bar for overview image = 100 µm; ROI1, scale bars = 50 µm). ROI2: zoom-in of age-associated B cells within the tertiary lymphoid-like structure. ROI3: Gzmk⁺ T cells located in alveolar septa adjacent to SFTPC⁺ alveolar type II (AT2) cells. Scale bars = 10 µm. n = 3 biological replicates.

Aging of the immune system is a key hallmark of organismal aging, leading to a state of immune dysfunction characterised by significant changes in innate and adaptive immune cell populations and chronic low-grade inflammation (inflammaging). Similarly, in the lung, we observed extensive immune cell remodelling with aging, evident already at baseline (0 dpi) and persisting after injury (Fig. 4E**-F**, Supplementary Fig. 11A-E). This included age-driven shifts in monocyte/macrophage and lymphocyte populations.

Among the most striking age-associated alterations in the myeloid compartment, we identified a depletion of Arginase 1⁺ (Arg1⁺) monocytes in aged lungs (Fig. 4E**-F**, Supplementary Fig. 11F-I). This monocyte population exhibited an activated phenotype, expressing markers of hypoxia (*Hif1a*) and pro-fibrotic factors (*Thbs1, Tgfbi, Tgm2, Vim*), alongside classical monocyte markers (*Ccr2, Vcan*) (Supplementary Fig. 11G). In young lungs, Arg1⁺ monocytes appeared in a time-dependent manner, emerging at 3 dpi, peaking during the fibrogenic phase, and returning to baseline by 37 dpi. However, this population failed to emerge in aged lungs following injury, suggesting a fundamental defect in their recruitment and/or activation during the regenerative response (Supplementary Fig. 11F).

Notably, a decline in Arg1⁺ monocytes/macrophages has recently been identified as a conserved hallmark of age-related regenerative decline across multiple tissues beyond the lung^36^, supporting its potential contribution to age-defective repair mechanisms.

Beyond myeloid cells, aging led to profound compositional shifts in the lymphocyte compartment. Aged mice exhibited a reduction in naïve T and B cells, coupled with an increase in memory T and B cells (Fig. 4E**-G**). In addition, aged lungs showed a substantial expansion of plasma cells, indicating increased antibody production and a heightened chronic immune activation state. Aging was further characterized by the emergence of distinct, age-specific T and B cell populations that exhibited unique transcriptional signatures compared to their conventional lymphocyte counterparts (Supplementary Fig. 11J). Relative frequency analysis showed an increase already at baseline in aged lungs that was largely unaffected by bleomycin challenge (Fig. 4G**-I**).

Among these, we identified a transcriptionally distinct subset of age-associated B cells. Compared to other B cell subtypes, these differentially expressed genes encoding surface receptors (*Fcgr2b, Cd22, Cd38, Cd81, Cd2*), transcription factors (*Tbx21, Zeb2, Zbtb32*), MHC signaling molecules (*H2-D1, H2-Q7I, H2-Ab1*), and metabolic regulators (*Pld4, Lgals1*). Conversely, canonical B cell markers such as *Cr2, Fcer2a*, and *Sell* were significantly downregulated, reinforcing their distinct molecular identity (Fig. 4J**, Supplementary** Fig. 11J). Functionally, age-associated B cells have previously been described as atypical memory B cells, which are enriched in autoantibodies and poised for plasma cell differentiation^37,38^. These cells have been shown to arise with aging and infections and are expanded in autoimmune diseases such as multiple sclerosis, systemic lupus erythematosus and rheumatoid arthritis, suggesting they may contribute to auto-inflammatory circuits in aged lungs^39^.

In parallel, we identified a transcriptionally unique population of age-associated CD8⁺ T cells marked by high expression of *Granzyme K (Gzmk)*. These Gzmk⁺ T cells co-expressed markers of tissue residency (*Itga4/CD49d, Cxcr6*), effector function (*Gzma, Gzmb*), and T cell exhaustion, including elevated levels of *Pdcd1 (PD-1)* and the transcription factor *Tox*, consistent with impaired classical cytotoxic activity (Fig. 4K**, Supplementary** Fig. 11J). This population has previously been shown to accumulate with age across multiple organs—including blood, spleen, liver, adipose tissue, and lungs, where they can constitute up to half of all CD8⁺ T cells^7^. While their relative abundance was not significantly altered by bleomycin challenge, injury modulated their phenotype (Supplementary Fig. 11K**/L**). Specifically, Gzmk⁺ T cells exhibited reduced expression of cytotoxic mediators (*Gzma, Gzmb, Prf1*), alongside increased expression of tissue-homing and activation markers (*Itgb1, Cxcr3*). These cells also showed enhanced activation of pro-inflammatory signaling pathways, including *Il6, Tnf,* and *Ifng*, as well as pathways related to epithelial cell differentiation (Supplementary Fig. 11K**/L**), suggesting a shift toward a tissue-remodeling, pro-inflammatory state that may exacerbate local inflammation.

To further characterize these populations, we examined their spatial distribution using multiplexed immunofluorescence. This confirmed the selective accumulation of age-associated B cells (*CD19*⁺ *T-bet*⁺) and Gzmk⁺ T cells (*CD8*⁺ *PD-1*⁺) in aged lungs (Fig. 4L), with absence in young lungs (Supplementary Fig. 11M). Both cell types were predominantly localized within tertiary lymphoid structures, composed of aggregates of B, T, and lymphatic cells that were situated near pulmonary veins and bronchioles. Additionally, Gzmk⁺ T cells were detected also rarely throughout the lung parenchyma, where they were observed near type II alveolar epithelial cells (AT2, Fig. 4L), suggesting potential functional interactions between the two cell types.

### Gzmk**⁺** T cells impair alveolar regeneration via induction of epithelial senescence

To functionally test this putative crosstalk between Gzmk⁺ T cells and AT2 cells, we employed an alveolar organoid system^28,40^ and extended it by incorporating T cells (Fig. 5A). To model the age-associated Gzmk⁺ T cell phenotype in vitro, we isolated T cells from the spleens of young mice and differentiated them into Gzmk+ T cells. IL-15 is an important cytokine driving inflammaging^41^ and induces the Gzmk⁺ T cell program^42,43^. Consistent with these previous studies, IL-15 treatment efficiently induced a Gzmk⁺ phenotype in vitro (Supplementary Fig. 12A**/B**). Notably, *Il15* expression was also significantly upregulated in aged lungs at baseline and following injury, paralleling increased *Gzmk* expression in lymphoid cells (Supplementary Fig. 12 C-E).

**Figure 5.**
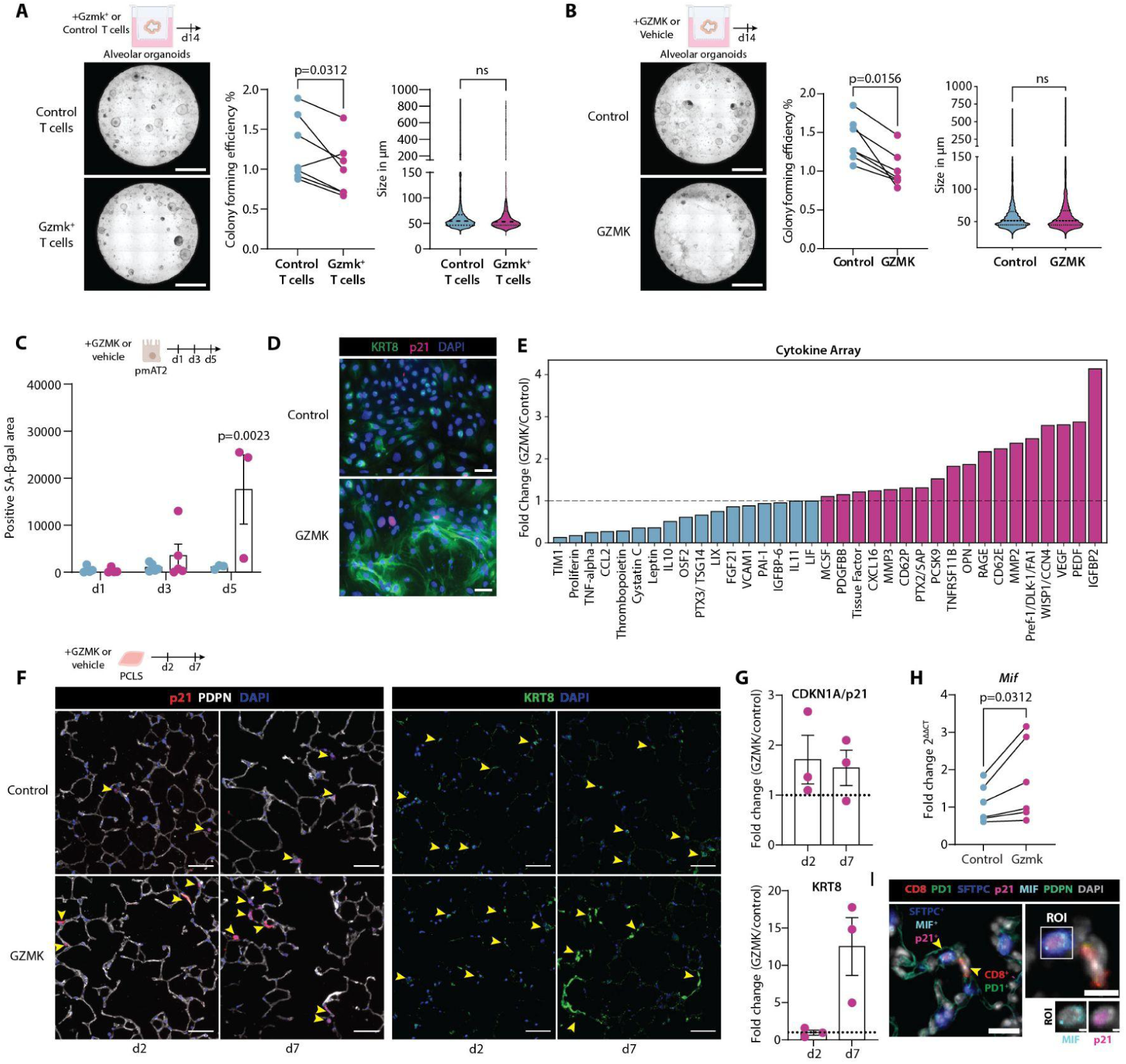
Gzmk⁺ T Cells Impair Alveolar Regeneration via Induction of Epithelial Senescence. (A, B) Colony forming efficiency (%) and individual organoid sizes (µm) of alveolar organoid cultures derived from young mice co-cultured with either IL-15-induced Gzmk⁺ or control T cells (A) or stimulated with 100 ng/mL recombinant GZMK or vehicle (B). Representative brightfield images of organoid cultures are shown. Scale bars = 50 µm. Statistics: Paired Wilcoxon test; n = 6–7 biological replicates. (C) Time-resolved analysis of senescence-associated β-galactosidase (SA-β-Gal)-positive stain area in 2D-cultured primary murine AT2 cells (pmAT2) stimulated with recombinant GZMK or vehicle. Statistics: Two-way ANOVA with Šídák’s multiple comparisons test; n = 3–5 biological replicates. (D) Representative immunofluorescence staining of KRT8 (green) and CDKN1A/p21 (pink) in pmAT2 cells stimulated with GZMK or vehicle at day 5. Scale bars = 50 µm. (E) Cytokine array of supernatants from pmAT2 cells stimulated with GZMK or vehicle at day 3. Ranked relative fold change (GZMK/Control) in protein expression is shown. Pool of 5 biological replicates. (F) Representative immunofluorescence images showing expression of CDKN1A/p21 (red), Podoplanin (PDPN, white), and KRT8 (green) in PCLS stimulated with 100 ng/ml recombinant GZMK or vehicle at day 2 or 7 post-stimulation. Scale bars = 50 µm; n = 3 biological replicates. (G) Quantification of immunofluorescence stainings shown in (F). Relative fold change (GZMK/Control) is presented. (H) Relative fold change in *Mif* mRNA expression in pmAT2 cells following stimulation with GZMK or vehicle at day 3. Statistics: Paired Wilcoxon test; n = 6 biological replicates. (I) Representative 4i image of lung tissue from aged mice showing CD8^+^/PD1^+^ Gzmk⁺ T cells adjacent to SFTPC^+^ AT2 cells positive for p21 and MIF expression. Scale bars = 20 µm for overview image, 10 µm for zoom-in and 2 µm for the selected ROI. n = 3 biological replicates.

When IL-15–induced Gzmk⁺ T cells were added to alveolar organoid cultures, we observed a significant reduction in colony-forming efficiency, yet not in organoid size compared to co-cultures with non-IL-15-treated T cells (Fig. 5A). This effect was not accompanied by increased cytotoxicity, suggesting a non-lytic mechanism of epithelial impairment (Supplementary Fig. 12F). To determine whether this impairment was mediated by soluble factors, we treated organoid cultures directly with recombinant GZMK. GZMK alone was sufficient to recapitulate the inhibitory effect on colony formation (Fig. 5B), indicating that it acts as a key effector molecule in this context. Importantly, the effect of GZMK was most pronounced when administered early during organoid development, suggesting that GZMK interferes with epithelial differentiation or early lineage commitment (Supplementary Fig. 12G).

Previous studies have shown that Gzmk⁺ T cells can promote senescence and SASP secretion in neighbouring cells, including fibroblasts^7^. We therefore investigated next whether GZMK could modulate the secretome and senescence phenotype of AT2 cells. Treatment of primary mouse AT2 cells with recombinant GZMK induced a time-dependent increase in senescence-associated β-galactosidase (SA-β-Gal) activity (Fig. 5C) and upregulation of CDKN1A/p21 in 2D-cultured primary AT2 cells (Fig. 5D) with no induction of cytotoxicity (Supplementary Fig. 12H). Consistently, GZMK treatment led to a pronounced shift in their secretory profile toward a SASP-like state, including upregulation of IGFBP2, VEGF, and TNFRSF11B (Fig. 5E). This was accompanied by an increased expression of KRT8 (Fig. 5D), indicative of a shift toward the Krt8-ADI state.

To validate these findings in a more physiologically relevant and complex model, we used precision-cut lung slices (PCLS) from young mice, which preserve the native lung tissue architecture and multicellular interactions. Consistent with our in vitro findings, GZMK treatment in PCLS induced upregulation of CDKN1A/p21 and KRT8 protein expression (Fig. 5F**/G)**. A similar effect was observed upon addition of in vitro–differentiated IL-15–induced Gzmk⁺ T cells into young PCLS (Supplementary Fig. 12I**/J**), confirming that Gzmk⁺ T cells can drive epithelial senescence and dysfunction.

Having previously observed elevated *Mif* expression in p21⁺ Krt8-ADI cells in aged mice in vivo, we next tested whether GZMK could directly induce *Mif* expression in epithelial cells. Indeed, GZMK stimulation significantly upregulated *Mif* mRNA levels in cultured primary AT2 cells (Fig. 5H). This was further validated in situ, where Gzmk⁺ T cells were found adjacent to AT2 cells co-expressing p21 and MIF (Fig. 5I), suggesting that Gzmk^+^ T cells are a predisposing driver of epithelial cell dysfunction, promoting accumulation of senescent Krt8-ADI which in turn sustain fibrosis via MIF-driven myofibroblast activation in aged mouse lungs.

## Discussion

Our longitudinal single-cell atlas provides a high-resolution view of how aging fundamentally alters the dynamics of lung regeneration following bleomycin-induced injury, shifting from a coordinated, transient repair process in young mice to a protracted, maladaptive response in aged ones that culminates in persistent fibrosis. We computationally analyzed multicellular programs associated with aging and identified age dependent shifts in the generation and resolution of injury-associated regenerative cell states (RCSs). For selected mediators of cellular crosstalk in aged mice we provide experimental evidence pointing to a specific epithelial-mesenchymal-immune crosstalk at the core of persistent fibrosis in old mice. To make our data easily accessible we provide an interactive data browser at https://hschillerlabshiny.shinyapps.io/BleomycinAging. Several aspects of our findings may extend to other organ systems. Persistent RCSs, including epithelial intermediates, activated fibroblasts, and inflammatory endothelial states, have also been observed in tissue injury such as the liver, kidney, and intestine following injury^44–46^. It is possible that mechanisms of immune-mediated impairment of stem cell niches, as shown here for the lung, may act in a similar way across organ systems^7,47^.

Although, we found induction of transitional cell states across multiple lineages in both age groups, their early-stage expansion was significantly attenuated in aged mice, suggesting delayed induction of repair processes. Furthermore, aged mice exhibited persistence of a subset of transitional cell states, all of which are also elevated in PF patients^11,14,19,48–50^. To which extent these shifts in temporal dynamics are driven by cell autonomous or cell-non-autonomous aging effects is unclear. On the one hand, we found big differences in stem cell intrinsic stress response pathways (e.g. DNA damage repair responses and gamma-H2ax foci), suggesting that to some extent cell autonomous roadblocks might cause the persistence of certain cell states (e.g. Krt8-ADI). On the other hand, we have shown that the aged immune niche, exemplified by the Gzmk-high CD8^+^ T cells, can exert non-cell autonomous effects on stem cell differentiation. Therefore, it is likely that a combination of these changes impairs regenerative responses and targeting both may be necessary for full rejuvenation.

As shown here and in similar work, aging is associated with the expansion of distinct immune cell subsets, including functionally altered B and T cell populations^4^. Although this immunological aging is systemic, certain organs, including the lung, appear to be more susceptible to early and severe alterations^51,52^. Consistent with our findings here, evidence suggests that immune cells can actively impair the regenerative capacity of epithelial cells in the lung^40,53,54^, thereby contributing to failed repair and CLD following injury. We found that increased secretion of GZMK in aged mice alone could be sufficient to shift the balance of pro-fibrotic cell states in the tissue. While classical CD8⁺ T cells exert cytotoxic effects via perforin and Gzmb, the Gzmk⁺ T cell subset we describe expresses low levels of canonical cytolytic genes but induces a non-cytotoxic, senescence-like response in alveolar epithelial cells via GZMK. Even though GZMK has been described as non-cytotoxic but pro-inflammatory in multiple studies^55^, our finding that GZMK directly impairs lung stem cell function is novel. We currently do not know the exact molecular mechanism of this effect on stem cells, but prior knowledge would suggest that GZMK enzymatic activity could modify the extracellular space and thereby specific signaling pathways. For instance, it has been shown that extracellular GZMK can directly regulate the complement cascade^8^ and PAR signaling^56^, which in turn have also been implicated in the modulation of cellular senescence^57^. Thus, our findings support an emerging concept that Gzmk⁺ T cells function as mediators of tissue remodeling rather than killers, particularly in the context of aging^7^. Also in a human disease setting, such as in the bronchoalveolar lavage of fibrotic hypersensitivity pneumonitis patients and in nasal tissue of patients with chronic rhinusitis, recent reports have described an increase of Gzmk⁺ T cells^8,58^.

We show that experimental GZMK treatment in precision cut lung slices induce pro-fibrogenic Krt8-ADI. In aged mice, we observed a strongly increased crosstalk of Krt8-ADI with Cthrc1⁺ myofibroblasts in the resolution phase of repair. MIF, which is a well described protein in the SASP^34,35^, emerged as an effector molecule increased by aging in this crosstalk, which we could validate functionally using in vitro experiments. Notably, this supports data showing that targeting MIF in experimental fibrosis reduced mortality and acute lung injury in bleomycin-treated mice^59^. In addition to MIF, our ligand-receptor modeling revealed a broader repertoire of profibrotic signals enriched in aged lungs. These included AREG, EREG, PTGS2, and SAA3, among others, which were increased in aged epithelial populations and predicted to activate myofibroblasts. Several of these ligands are known components of the SASP or are involved in extracellular matrix remodeling, angiogenesis, and immune activation^32,33,60^. In particular, the sustained expression of amphiregulin (Areg) in Krt8-ADI has recently been shown to drive progressive fibrosis in mice^60^. It seems that Krt8-ADI can effectuate fibrosis via multiple secreted ligands and therefore combination strategies or interference more upstream may be required to disrupt the broader fibrogenic signaling network.

We acknowledge several limitations of our study. We do not know how long the described pro-fibrotic circuit of cells can persist, as the current design has not addressed truly long lasting consequences beyond 37 dpi. A recent report on transitional cell states post influenza infection suggests however that such injury induced cell states can persist indefinitely^19^. Since we did address the role of Gzmk⁺ T cells in ex vivo organoid assays rather than in vivo we do not currently know to what extent the removal of these cells would lead to a phenotypic rescue in old mice. Future studies might explore causal interventions with specific depletion of the Gzmk⁺ CD8 T cells or targeting Gzmk itself in aged models, to dissect this axis and potentially translate into anti-fibrotic therapies.

In conclusion, we provide a high resolution and time-resolved single cell atlas of lung repair in the context of aging and use single-cell interpretable tensor decomposition to offer a novel perspective, identifying strongest aging associations with T/B lymphocytes and macrophages, thus challenging epithelium-centric views of lung aging and repair and emphasizing multicellular coordination. These insights provide a framework for developing therapeutic strategies that restore regenerative competence in aged lungs by modulating immune niches, or interrupting specific crosstalk signals such as GZMK or MIF.

## Methods

### Animal experiments

Pathogen-free, young (12-weeks-old) and aged (18-months-old) female and male C57BL/6J-rj mice were purchased from Janvier-Labs (Le Genest-Saint-Isle, France). Mice were randomized into control (untreated/day 0) or bleomycin-treated groups, and lung injury and fibrosis was induced by an oropharyngeal, single-dose instillation of 2U/kg bleomycin sulphate (Sigma Aldrich, Germany). Mice were sacrificed at days 0, 3, 10, 20, 30 and 37 post-bleomycin injury. All animal experiments were conducted in strict accordance with governmental and international guidelines for animal research and were approved by the local government of Upper Bavaria (license number: ROB-55.2-2532.Vet_02-16-208).

### Generation of single-cell suspensions from murine lung tissue

Single cell suspensions from lung tissues were generated as described in *Strunz, Simon et al.*^11^. In brief, mouse lungs were transcardially perfused with sterile PBS, and the left lungs were inflated with an enzymatic mix composed of dispase (50 units/ml), collagenase (156 Units/ml, elastase (3.125 units/ml), and DNase (17 units/ml), minced, and enzymatically digested for 20 min at 37 °C with constant shaking. Enzymatic activity was inactivated by adding PBS supplemented with 10% fetal calf serum (FCS). The dissociated cells were passed through a 70 µm cell strainer, pelleted by centrifugation (300xg, 5 minutes), and subjected to red blood cell lysis for 1 minute. The lysis reaction was stopped with PBS containing 10% FCS, followed by another centrifugation step (300xg, 5 minutes). The resulting cell pellet was resuspended in PBS with 10% FCS, assessed for viability, and manually counted. For Dropseq-based scRNA-seq, cells were aliquoted in PBS supplemented with 0.04% of bovine serum albumin at a final concentration of 150 cells/µl.

### Dropseq-based scRNA-seq

Single-cell profiling using Drop-seq was performed as previously described^11^. In brief, single cells (150 cells/µl) were co-encapsulated in droplets with barcoded beads (120 beads/µl, ChemGenes Corporation, Wilmington, MA) using an in-house fabricated microfluidic device at a flow rate of 4000 µl/h. Droplet emulsions were collected for 20 minutes per sample, followed by droplet breakage. Beads were harvested, and hybridized mRNA transcripts were reverse transcribed using Maxima Reverse Transcriptase (Thermo Fisher) with a template-switch oligonucleotide primer (50 μM; sequence: AAGCAGTGGTATCAACGCAGAGTGAATrGrGrG). Unused primers were removed by exonuclease I (New England Biolabs), followed by second-strand synthesis using the Klenow enzyme. Beads were then washed, counted, and aliquoted for pre-amplification. PCR products from each sample were pooled, purified, and quantified using the BioAnalyzer High Sensitivity DNA Assay (Agilent). For each sample, 1 ng of pre-amplified cDNA, corresponding to an estimated 3000 cells, was tagmented using Nextera XT (Illumina) with a custom P5 primer (10 μM; Integrated DNA Technologies; sequence: AATGATACGGCGACCACCGAGATCTACACGCCTGTCCGCGGAAGCAGTGGTATCAACGCAGAGT*A*C). The final single-cell libraries were sequenced on a NovaSeq 6000 instrument.

### Preprocessing and analysis of scRNA-seq data

For scRNA-seq processing the Drop-seq computational pipeline (version 2.0) was used, as described by *Macosko et al.*^61^. Briefly, read alignment was performed using STAR (version 2.5.2a) against the mm10 reference genome. During quality control (QC), genes with fewer than one count and those expressed in fewer than three cells were removed. Additionally, cells with 10% or more mitochondrial counts, fewer than 200 detected genes, or fewer than 200 total counts were excluded. To further refine the dataset, doublet detection was carried out at the individual sample level using Scrublet^62^. Ambient RNA contamination was corrected with SoupX^63^, where the background contamination fraction was manually set to 0.3. Normalization was performed using scran’s size factor-based approach, with cell groupings obtained through Louvain clustering (resolution = 2.0) as input. Subsequently, log transformation was applied using Scanpy’s pp.log1p() function. Highly variable genes (HVGs) were identified separately for each sample, selecting the top 6000 genes per sample. Genes were considered as overall HVGs only if they were highly variable in at least 16 samples. Principal Component Analysis (PCA) was performed using the HVGs as input, followed by the construction of a k-nearest neighbors (kNN) graph based on the first 50 principal components. Graph-based clustering was then performed using the Leiden algorithm, and clusters were annotated based on canonical marker genes. Initially, a coarse-grained annotation was performed to classify cells into four major subsets: epithelial cells (Epcam+), stromal cells (Col1a1+), endothelial cells (Cldn5+), and immune cells (Ptprc+). Subsequently, HVG selection, PCA, and Leiden-based graph-based clustering were repeated within each subset to achieve a more fine-grained annotation. Marker genes were identified using a Wilcoxon rank-sum test, with genes considered as markers if the false discovery rate (FDR)-corrected p-value was below 0.05.

### Cell perturbation analysis with Augur

To assess how cell types respond to bleomycin-induced injury in young and old mice, we performed Augur analysis^64^ as implemented in the Python package pertpy (https://doi.org/10.1101/2024.08.04.606516) (version 0.7.0), using the following parameters: n_subsamples=100, subsample_size=20, select_variance_features=True.

### Differential abundance testing with Milo

Milo^18^ was applied to perform differential abundance testing by assigning cells to overlapping neighborhoods within a k-nearest neighbor graph. Milo was run with default parameters as implemented in the pertpy Python package.

### Differential gene expression (DGE) analysis

Differential gene expression (DGE) analysis was performed using diffxpy (version 0.7.4) as previously described^65^. We performed a Wald test with default parameters for all genes that were expressed in at least 10 cells. We considered genes differentially expressed at a statistically significant level if the FDR-corrected p-value<0.05.

### Clustering of temporal gene expression profiles

DGEs were identified by comparing young and aged mice at day 37 post-injury using pseudobulk expression profiles using diffxpy. Genes were considered significantly differentially expressed if they met a FDR-adjusted p-value< 0.05. Upregulated genes were defined by a log₂ fold change ≥ 0.25, and downregulated genes by a log₂ fold change ≤ –0.25. To enable comparison across time points, the average expression profiles of the selected DEGs were z-score normalized separately for the young and aged cohorts. To identify groups of genes with similar temporal expression dynamics, a Fréchet distance correlation matrix was computed using the z-scored expression profiles from the aged cohort (via the similaritymeasures Python package). Agglomerative hierarchical clustering was applied to the precomputed correlation matrix to group genes with shared expression trajectories. The clustering algorithm used complete linkage and was implemented using the scikit-learn Python package. Cluster-specific gene sets were extracted for downstream analysis. To visualize the differences in gene expression trajectories, the z-scored, time-resolved expression data of each gene cluster were plotted separately for both old and young mice cohorts. Two types of visualizations were generated: expression heatmaps and expression-averaged line plots, providing a detailed comparison of gene expression dynamics between both age groups.

### Gene set enrichment analysis of temporal gene clusters

To functionally characterize each gene cluster, the corresponding genes were subjected to gene set enrichment analysis. The analysis was performed using the “GO_Biological_Process_2023” and “MSigDB_Hallmark_2020” databases for *Mus musculus*, utilizing the GSEApy tool with default parameters. A significance threshold of FDR-adjusted p-value ≤ 0.05 was applied to identify enriched biological processes and hallmark pathways. The results of the enrichment analysis were visualized using heatmaps. For each cluster, enriched terms from the “MSigDB_Hallmark_2020” and “GO_Biological_Process_2023 database were manually pre-selected to reduce figure complexity. These terms were visualized based on their -log10 p-value, with heatmap visualizations generated using the R pheatmap package.

### Pathway scoring for temporal gene clusters

To evaluate time-resolved pathway activity for the terms enriched within the gene clusters sharing similar expression trajectories, the complete gene lists corresponding to each pathway identifier were retrieved from the MSigDB_Hallmark_2020 and GO_Biological_Process_2023 databases. Pathway scores were calculated as the average expression of the gene set subtracted by the average expression of a reference gene set. This calculation was performed using the score_genes function of the scanpy Python package with default parameters. The resulting pathway scores were visualized using the matrixplot function of scanpy, enabling a comprehensive comparison of pathway activities over time.

### Senescence gene signature scorings

Gene signature scores for cell senescence (SenMayo^34^, FRIDMAN_SENESCENCE^66^, SENESCENCE (Strunz et al.)^11^, SASP^35^) and related biological processes (HALLMARK_P53_PATHWAY, HALLMARK_DNA_REPAIR, HALLMARK_INFLAMMATORY_RESPONSE, HALLMARK_PROTEIN_SECRETION) were retrieved from the respective publications or MSigDB database. The respective scores were using Scanpy’s sc.tl.score_genes() function with default parameters.

### Cell-cell communication analysis

Cell-cell communication analysis was conducted using the R package *NicheNet*^67^ as previously described^65^. To define perturbed gene programs in young and old mice following bleomycin injury, we performed DGE analysis for each potential receiver cell type. Genes were included in the perturbed gene programs if they met the following criteria: an FDR-corrected p-value < 0.05, an absolute log_2_ fold change > 0.5, and expression in at least 10% of the receiver cells in the perturbed condition. These perturbed gene programs were then used as input for *NicheNet* to predict ligands that could induce the gene program in the receiver cell types.

### ScITD

We applied the scITD (single-cell Interpretable Tensor Decomposition) framework^17^ using the official R implementation (v1.0.4) to analyze single-cell RNA-seq data from aging mice. Cell types with low abundance (Megakaryocytes, Pericytes, SMCs, and NK cells) were excluded. A gene expression tensor was constructed by averaging normalized expression values per gene and cell type across samples, resulting in a 3-mode tensor with dimensions: donors × genes × cell types. Tensor construction (form_tensor) included normalization (regular method, scale factor = 10,000), selection of highly variable genes based on normalized variance p-values (threshold = 0.01), and variance scaling. We performed Tucker decomposition using run_tucker_ica with ranks set to (5,16), extracting five multicellular programs that vary across donors. Donor factor scores were statistically associated with metadata variables (treatment, age, day) using p-values. To quantify how individual cell types contributed to each factor, we extracted the gene loadings from the second tensor mode, parsed the cell-type labels, and calculated the mean absolute loading per cell type and factor.

### Primary mouse AT2 cells 2D culture

Female C57BL6/J mice were euthanized by an intraperitoneal overdose of ketamine and xylazine. Lungs were then flushed via the right ventricle with PBS and filled with Dispase (Corning Incorporated USA). Filled lungs were excised and further digested in Dispase for 40 minutes at room temperature. The lungs were then mechanically minced with forceps and suspension was sequentially filtered through 100µm (Corning Incorporated, USA, 431752) and 40µm cell strainers (Corning Incorporated, USA, 431750) to obtain a single cell suspension. This suspension was then incubated in petri plates for 30 mins to deplete fibroblasts via adhesion. This suspension was then depleted of immune and endothelial cells via MACS separation using anti-CD45 (Miltenyi Biotec, Germany) and anti-CD31 magnetic microbeads (Miltenyi Biotec, Germany, 130-052-301), respectively, according to manufacturer’s instructions. Finally, 1.5-2.0×10^6^ cells (6-well) or 3.7-3.8 ×10^5^ (24-well) were seeded in 2 ml or 0.5ml medium, respectively. The cells were cultured in a medium with the following composition: DMEM/F-12 supplemented with 3.6 mg/ml D-Glucose, 2% GlutaMAX™, 1% Penicillin/Streptomycin, 10 mM HEPES, 10%. FCS. 48h after seeding, cells were treated with vehicle control or medium containing 100ng/ml mouse GZMK (RPB209Mu01, Cloud-Clone). After 24h and 72h, the supernatants were collected and frozen and the cells were either fixed with 4% PFA for immunofluorescence or frozen directly for RNA isolation at −80°C.

### Isolation of AT2 cells for organoid cultures

Female C57BL6/J mice were euthanized by an intraperitoneal overdose of narcotics. Lungs were then perfused via the right ventricle with PBS and filled with Dispase (Corning Incorporated USA, 354235). Filled lungs were excised and further digested in Dispase for 40 minutes at room temperature. The lungs were then mechanically minced with forceps and suspension was sequentially filtered through 100 µm (Corning Incorporated, USA, 431752) and 40 µm cell strainers (Corning Incorporated, USA, 431750) to obtain a single cell suspension. This suspension was then depleted of immune and endothelial cells via MACS separation using anti-CD45 (Miltenyi Biotec, Germany) and anti-CD31 magnetic microbeads (Miltenyi Biotec, Germany, 130-052-301), respectively, according to manufacturer’s instructions. This depleted fraction was further enriched for distal lung epithelial progenitors via MACS separation using anti-CD326 EpCAM magnetic microbeads (Miltenyi Biotec, Germany, 130-105-958). This cell suspension was kept on ice until further use.

### Isolation and differentiation of splenic T cells

Isolation and differentiation of splenic T cells was performed as previously described^42^. One million per ml splenic CD8^+^ T cells from C57BL/6 mice purified by immunomagnetic separation using AutoMACS (Miltenyi Biotec) were activated with plate-bound anti-CD3 (2 μg/ml ml) in RPMI 1640 medium containing FCS (10%), L-glutamine (2 mM), penicillin–streptomycin (100 U/ml) and IL-2 (100 U/ml) for 2 days in 24-well plates. Cells were then removed and transferred to a new plate at a concentration of 10^6^ cells per ml in the presence of TGF-β1 (5 ng/ml, ThermoFisher, no. 100-21). After 2 days, vital cells were isolated through Pancoll density centrifugation (1,440xg, 20 min). Vital CD8+ T cells were stimulated with IL-15 (10 ng/ml, ThermoFisher, no. 200-15) for 24 h to become highly enriched for Gzmk+ T-cells^42^.

### High throughput Fibroblast into Myofibroblast Transition (FMT) assay

The study was approved by the local ethics committee of the Ludwig-Maximilians University of Munich, Germany (Ethic vote 19-630). Written informed consent was obtained for all study participants. Primary human lung fibroblasts (phLF) were obtained from the CPC-M bioArchive at the Comprehensive Pneumology Center (CPC Munich, Germany). The isolation of primary human lung fibroblasts (phLF) from IPF lung tissue was carried out as previously described^30^. IPF fibroblasts were seeded in a 96-well plate (15 x 10^3^ cells/ well) in DMEM/F-12 (Life Technologies, UK, 11320033) supplemented with 20 % FBS (Pan Biotech, Germany, P30-3702) and 1% Penicillin/Streptomycin (Life Technologies, UK, 15140122) for 24 hrs. The following day, cells were synchronized in serum-starved media (DMEM-F12, 1% FBS, 1% Penicillin/Streptomycin) for 24 hrs before treatment. Subsequently, the cells were stimulated with vehicle control (0.1%BSA), or recombinant MIF (100 ng/ml, R&D systems, USA,) in starvation media for 48 hrs. The stimulated cells were fixed with cold methanol for 5 minutes and stained overnight with α-SMA (Sigma-Aldrich, Germany, A5228, 1/200) and DAPI (Sigma-Aldrich, Germany, D9542, 1/1000). Staining was continued with Alexa Fluor 568 (Invitrogen, Germany, A10037) as a secondary antibody for 2 hours at room temperature in the dark. Then, the middle part of each well was imaged with the fluorescence microscope AxioImager (Zeiss, Germany) at 20X magnification. Finally, images were analysed using a custom Jupyter Notebook pipeline that automated the mean fluorescence intensity (MFI) of α-SMA+ cells normalized by the number of nuclei.

### Alveolar organoid cultures

5×10^5^ CD31^-^CD45^-^CD326^+^ lung cells were combined with 5×10^5^ CCL206 fibroblasts (ATCC, MLg 2908) that have been growth inhibited using Mitomycin C (MP Biochemicals, USA, 02100498-CF). For T cell co-culture experiments, isolated unstimulated and IL-15 stimulated T cells were mixed in a 1:4 ratio with the cell mix described before. In both cases, cell solutions were further mixed 1:1 with Growth Factor Reduced Matrigel (Corning Incorporated, USA, 354230), seeded in 96-well plates and allowed to solidify for 15 mins at 37°C 5%CO_2_. Cells were then cultured in DMEM/F-12 (Life Technologies, UK, 11320033) supplemented with 5% FBS, 1X Penicillin/Streptomycin (Life Technologies, UK, 15140122), 1X Amphotericin B (Life Technologies, UK, 15290026), 1X GlutaMAX (Life Technologies, UK, 35050061), 1X Insulin-Transferrin-Selenium-Sodium Pyruvate (Life Technologies, UK, 51300044), 0.1µg/ml Cholera Toxin (Merck-Sigma-Aldrich, Germany, C8052), 0.025µg/ml of recombinant Mouse Epidermal Growth Factor (Merck-Sigma-Aldrich, Germany, SRP3196), 30µg/ml Bovine Pituitary Extract (Sigma-Aldrich, Germany, P1476), and 0.01µM Retinoic Acid (Merck-Sigma-Aldrich, Germany, 223018) for 14 days. The Rock inhibitor (10µM, Y-27632 dihydrochloride (Merck-Sigma-Aldrich, Germany, Y0503)) was added only for the first 48h. Organoids were treated with 100ng/ml recombinant mouse Granzyme K (Cloud-Clone Corp., RPB209Mu01) and 5ng/ml recombinant MIF (R&D Systems, USA) every 48h until day 14. The culture supernatants were collected, and organoids were washed once in DPBS. Then, Organoids were imaged on a Carl Zeiss Axio Imager using Brightfield (50X magnification). Finally, organoids were quantified and sized using an in-house R-CNN based counter^68^.

### Generation of mPCLS

mPCLS were obtained from murine lungs pre-filled with 2% low gelling temperature agarose DMEM-F12 (Thermo Scientific, USA) and kept at 4°C for 30 min before slicing. 300 µm mPCLS were generated using a 7000smz-2 Vibratome (Campden Instruments, England) as previously described^69,70^. Slices were cultured as 4mm punches in 96-well plates. mPCLS were cultured for up to 7 days in DMEM-F12 supplemented with 0.1% FBS, 1%penicillin/streptomycin and 1% amphotericin B and the medium was changed every 48/72 hours.

### mPCLS treatment with granzyme K

Previously prepared 4mm mPCLS in 96 well plates were treated with 100ng/ml recombinant GZMK or vehicle in supplemented PCLS medium. At day 2 and 5, tissue samples for RT-qPCR and immunofluorescence as well as the supernatants were collected. Treatment with recombinant GZMK or vehicle was replenished after the initial 48 h.

### Coculture of mPCLS with CD8+ T cells

Previously prepared mPCLS samples in 96-well plates were seeded with either 100 μl of PCLS medium (control treatment), or 100 μl of unstimulated or IL-15 stimulated T cells (10 000 cells/well) in supplemented PCLS medium. At day 2 and 5, tissue samples for immunofluorescence were collected. Additional 100ul of fresh medium were added on top of mPCLS after 48h of culture.

### LDH Assay

LDH Cytotoxicity WST Assay (Enzo Life Sciences GmbH, Germany, ENZ-KIT157) was used to assess cytotoxicity in supernatants from organoids and pmAT2 cells. Lysis buffer (1:10) was added to the lysis control samples incubated at 37°C for 30 min and supernatant was collected until use. For the assay, 100µl of medium (negative control), lysis control, or supernatants were mixed with 100µl of working solution in a 96 well plate. This mixture was incubated at room temperature for 30 minutes. 50µl of stop solution was added to each well. Absorbance was measured at 490nm using a TECAN Sunrise(TM) microplate reader. Absorbance values were calculated after removal of background signal (only medium) and percentage of cytotoxicity was calculated based on lysis control (100% cell death).

### RNA isolation from cultured cells

For RNA isolation, the peqGOLD Total RNA Kit from VWR was used, according to the manufacturer’s instructions. Briefly, cells were lysed with TRK Lysis Buffer. The lysate was homogenized using an RNA homogenizer and then centrifuged. Ethanol (70%) was added to the lysate, which was then transferred to an RNA mini column. Samples were washed with RNA wash buffer I and centrifuged. DNase I digestion (Qiagen or VWR) was performed, and the cells were again washed with RNA wash buffer I. This was followed by two washes with 80% ethanol. The columns were dried, and the RNA was eluted with Nuclease-free Water. RNA concentration was measured using a Nanodrop1000 (PeqLab). Finally, the eluted RNA was stored at-80°C.

### RT-qPCR

cDNA synthesis and RT-qPCR 500-1000ng RNA were denatured in 20μl of RNAse-free water (15 minutes at 70°C). For cDNA synthesis, the reaction mix was prepared using the following components: 10 µM Random Hexamers (Invitrogen, Cat. No. N8080127), 2 mM dNTP mix (Thermo Scientific, Cat. No. R0192), 1× First-Strand Buffer (Invitrogen, Cat. No. 1805701), and 40 mM DTT (Invitrogen, Cat. No. 1805701). Reverse transcriptase (Invitrogen, Cat. No. 28025013) was added at a final concentration of 10 U/µl, and RNase inhibitor (Applied Biosystems, Cat. No. N8080119) was included at 4 U/µl. All reagents were mixed on ice according to the manufacturer’s instructions. Then, the cDNA reaction mix was added to each sample and incubated for one cycle at 20°C for 10 min, one cycle at 43°C for 75 min, and one cycle at 99°C for 5 min. Finally, cDNA was diluted with RNAse-free water and kept at −20°C until use. The RT-qPCR reaction mix was prepared using 1X Light Cycler 480 SYBR Green Master mix and the desired primer pair (final concentration 5μM). Samples were loaded in a 96-well plate and incubated in a LightCycler 480II (Roche, Germany) following these steps: Pre-incubation: (1 cycle) 50°C for 2 min. Denaturation: (1 cycle) 95°C for 5 min. Amplification: (45 cycles) at 95°C for 5s, 59°C for 5s, 72°C for 5s. Melting curve: (1 cycle) 95°C for 5s, 60°C for 1min. Cooling: (1 cycle) 40°C for 30s. A two-derivative analysis was used to determine Ct values and the 2–ΔΔCt method was used to calculate the relative fold gene expression to each control.

### Cytokine array

A cytokine array was used to characterize the SASP from AT2 cells treated or not with GZMK for 72 hours. For this, supernatants were concentrated using the Amicon Ultra-0.5 Centrifugal Filters 3 kDa MWCO (Merck, Germany), according to the manufactureŕs instructions using molecular biology grade water as exchange buffer. Total protein content was determined using the Pierce™ BCA Protein Assay Kit (Thermo Scientific, USA) for microplates. In short, 25 µl of sample or standard were added in duplicates and mixed with 200 µl of the working solution. The microplate was incubated for 30min at 37°C and absorbance was measured at 562 nm using a microplate reader Tecan Sunrise (Tecan GmbH, Germany). Absorbance values were then calculated by subtracting the background signal and protein concentration was calculated by interpolation of a linear regression using GraphPad Prism 9.5.1. Based on the protein concentration, supernatants from 5 biological replicates were pooled for a final concentration of 50ug of protein per sample. Then, manufacturer’s instructions (R&D Systems, USA) were followed for the cytokine array assay: Membranes were blocked using Array Buffer 4 for 1h at RT. Then, samples were diluted in Buffer 4 and 6 for a final volume of 1.5ml, added to each membrane, and incubated at 4°C overnight in a rocking platform.

The next day, membranes were washed 3 times with 1X wash buffer (10 min at RT, each time) and incubated with Detection Antibody Cocktail for 1h at RT in a rocking platform. Then, mem branes were washed 3 times with 1X wash buffer (10 min at RT, each time) and incubated with 1X Streptavidin-HRP for 30min at RT in a rocking platform. Finally, membranes were washed 3 times with 1X wash buffer (10 min at RT, each time), transferred to a plastic sheet protector, and the Chemi Reagent Mix was added to each membrane. Excess liquid was removed, membranes were incubated for 1 min and Chemiluminescence was detected using a ChemiDoc XRS+ Universal Hood II (Bio-Rad, USA). Then, the average pixel density from the duplicate spots was calculated after subtracting the background signal. The fold change (GZMK/CTRL) was quantified.

### Histology and Ashcroft scoring

For histology, perfused caudal lobes of the right mouse lung were inflated with 10% neutral-buffered formalin solution and fixed overnight at 4°C. After embedding in paraffin, lung sections were cut at a thickness of 3.5 μm and stained with hematoxylin and eosin (H&E) for analysis of the overall lung architecture and the presence of cellular infiltrates, and with Picrosirius Red to detect collagen deposition using standard protocols as previously described^71^. For semi-quantitative assessment of murine and human lung fibrosis, the Ashcroft Score was applied on Picrosirius Red stained lung sections as described previously^71^. Two blinded examiners (JGS, HW) performed the scorings in duplicates. If deviations of more than 1 score were observed, the respective slides were re-assessed to reach consensus.

### Immunofluorescence (IF) stainings, microscopy, and quantification of murine lung tissues and murine precision-cut lung slices

Formalin-fixed paraffin-embedded (FFPE) sections from bleomycin-treated and control mice, as well as murine precision-cut lung slices (mPCLS) were dried at 60°C for 30 min to 1 hour, followed by deparaffinization and rehydration through sequential incubations in xylene (2x 5 min), 100% ethanol (2×3 min), 90% ethanol (1×3 min), 80% ethanol (1×3 min), 70% ethanol (1×3 min), and Milli-Q water (1x 5 min). Heat-induced antigen retrieval was performed using 10 mM citrate buffer (pH 6.0) with one cycle at 125°C for 30 seconds followed by one cycle at 90°C for 10 seconds. After antigen retrieval, slides were washed twice with 1X PBS and blocked for 1 hour at room temperature with 10% Normal Donkey Serum in DAKO Antibody Diluent (Agilent Technologies, USA). When mouse anti-mouse primary antibodies were stained, an additional blocking step was performed using the M.O.M.® (Mouse on Mouse) Blocking Reagent (Vector Laboratories, USA) for 1 hour at room temperature. Subsequently, sections were incubated overnight at 4°C with primary antibodies (p21, abcam, ab188224, 1:800; KRT8, DSHB, TROMA-I, 1:200; PDPN, R&D, AF3670, 1:200; yH2Ax, Sigma, 05-636, 1:200) diluted in DAKO Antibody Diluent. The following day, samples were washed twice with 1X PBS and incubated with secondary antibodies (donkey anti-mouse AF488, Invitrogen, A21202, 1:250; donkey anti-rat AF488, Invitrogen, A21208, 1:250; donkey anti-rabbit AF568, Invitrogen, A10042, 1:250; donkey anti-goat AF647, Invitrogen, A21447, 1:250) diluted in 1% BSA/PBS for 2 hours at room temperature. After secondary antibody incubation, sections were counterstained with DAPI (2.5 µg/ml) for 10 minutes at room temperature and mounted using DAKO Fluorescence Mounting Medium (Agilent Technologies, USA). Imaging was performed using an AxioImager.M2 fluorescence microscope (Zeiss, Germany) with a 20× objective. For quantification, three to five randomly selected regions of interest were captured per sample. Image analysis was conducted using ImageJ, with nuclear proteins such as CDKN1A/p21 and γH2Ax quantified based on nuclear overlap with the DAPI signal, while mean fluorescence intensity (MFI) was normalized by the number of nuclei.

### Iterative indirect immunofluorescence imaging (4i)

Iterative indirect immunofluorescence imaging (4i) was performed as previously described^65^. Similar to the conventional immunofluorescence staining, FFPE sections were first deparaffinized, dehydrated, and then subjected to heat-mediated antigen retrieval using 1X R-universal antigen retrieval buffer (Aptum). To minimize nonspecific antibody binding, sections were blocked for 1 hour at room temperature (RT) with 1% BSA supplemented with 150 mM maleimide.

Primary antibodies were then applied and incubated overnight at 4°C. After thorough washing, secondary antibodies were added and incubated for 2 hours at RT, followed by DAPI staining and autofluorescence blocking. To prevent photocrosslinking during imaging, sections were immersed in an imaging buffer containing 700 mM N-acetylcysteine and 20% 1 M HEPES (pH 7.4) and imaged using the Axioscan 7 slide scanner (Zeiss). Antibody elution was performed using a stripping buffer (pH 2.5) containing 0.5 M L-glycine, 3 M urea, 3 M guanidinium chloride, and 70 mM TCEP-HCl. Subsequent rounds of antibody staining and imaging were conducted as described above, without repeating deparaffinization, dehydration, or antigen retrieval, until the desired plexity was achieved. The following primary (i) and secondary (ii) antibodies were used: i) p21 (abcam, ab188224, 1:800), KRT8 (DSHB, TROMA-I, 1:200), PDPN (R&D, AF3670, 1:200), MIF (abcam, ab65869, 1:500), TBET (Cell signaling, 97135, 1:100), CD3 (Sigma, C7930, 1:300), CD8a (Invitrogen, 14-0808-82, 1:500), LYVE-1 (R&D, AF2125, 1:500), pro-SPC (Millipore, AB3786, 1:200), PD1 (R&D, AF1021, 1:100), aSMA (Sigma, A5228, 1:1500), TRKB (R&D, AF1494, 1:40), PRX (Atlas antibodies, HPA001868, 1:500), CD19 (eBiosciences, 14-0194-82, 1:400). ii) donkey anti-mouse AF488 (Invitrogen, A21202, 1:250), donkey anti-rat AF488 (Invitrogen, A21208, 1:250), donkey anti-rabbit AF568 (Invitrogen, A10042, 1:250), donkey anti-goat AF647 (Invitrogen, A21447, 1:250). Alignment of microscopy images of different cycles was performed using an in-house generated software <u>ida-mdc / image-registration-tool ·</u> <u>GitLab</u> (https://github.com/schillerlab/2025_Aging_Bleo) by using the first DAPI cycle as reference.

### IF staining of cell cultures

The cells were fixed with 4%PFA at RT for 15 min and washed three times with PBS. Then, cells were permeabilized with 0.5% PBS + Triton-X for 15 min at RT. Then, samples were blocked with a 5% donkey serum solution in 0.1% PBST for 1 hour at RT and primary antibodies against P21 (abcam. ab188224, 1:800) and Krt8 (DSHB, TROMA-I, 1:200) were diluted in 1% donkey serum in 0.1% PBST, added to the samples and incubated overnight at 4°C. Next day, cells were washed 3 times with 0.1% PBST and then incubated with secondary antibodies (donkey-anti-rabbit AF568, Invitrogen A10042, donkey anti-rat AF488, Invitrogen, A21208) and DAPI diluted 1:500 in 1% donkey serum in 0.1% PBST for 2h at RT. The cells were again washed 3 times with 0.1% PBST. The coverslips were then mounted on glass slides using DAKO mounting medium and an AxioImager.M2 (Zeiss, Germany) was used to image the cells. Three regions of interest were captured. MFI and nuclei number were quantified using ImageJ software.

### Senescence Associated B-galactosidase staining

Primary AT2 cells were cultured and treated with GZMK as previously described. After 24h, 72h and 120h, the supernatant was removed and cells were washed once with 1X PBS. The cells were then stained for Senescence Associated B-galactosidase activity using the Senescence β-Galactosidase Staining Kit (Cell Signalling Technologies #9860) according to manufacturer’s Instructions. The stained cells were then imaged using a Brightfield Microscope with a phase contrast filter. The images were then analysed for β-Galactosidase levels using ImageJ.

### Statistics

All statistical analyses were conducted using GraphPad Prism (version 10.3.1) or the scipy.stats module in Python. For non-parametric unpaired comparisons between two groups, the Mann–Whitney U test was used. For comparisons across more than two unpaired groups, the Kruskal-Wallis test followed by Dunn’s multiple comparisons post hoc test was applied. For paired data, the Wilcoxon signed-rank test was used for two-group comparisons. In cases involving repeated measures across more than two conditions, two-way ANOVA with Šídák’s multiple comparisons correction was employed. All P-values less than 0.05 were considered statistically significant unless otherwise specified. When applicable, adjustments for multiple testing were applied to control the false discovery rate. \**p* < 0.05, \*\**P* < 0.01, \*\*\**p* < 0.001, and \*\*\*\**p* < 0.0001.

## Data availability

All data associated with this study are present in the paper and can be explored via the publicly available data visualization tool (https://hschillerlabshiny.shinyapps.io/BleomycinAging/). Raw data has been submitted to Sequence Read Archive (SRA) under the Bioproject (PRJNA1293810). Count Matrices are deposited at the Gene Expression Omnibus repository.

## Code availability

Code and Jupyter notebooks that were used to analyze the data are deposited at GitHub (https://github.com/schillerlab/2025_Aging_Bleo) and will be archived at Zenodo after publication.

## Author Contributions

J.G.S., M.L., and H.B.S. conceived and designed the study, supervised the overall project, and wrote the manuscript. J.G.S., C.H.M., I.A., and Y.C. conducted the animal work and performed the scRNA-seq experiments. J.G.S., Y.C., H.W., M.B., C.M., and K.W. performed and analyzed histological and immunostaining experiments. J.G.S. and H.B.S. led the analysis of the scRNA-seq data. J.G.S., M.A., D.L., A.M., S.P., S.K., and Sa.K. carried out the bioinformatic analyses of the scRNA-seq data. C.M., G.B., S.A., E.J., N.P., C.S., A.E., and M.D. performed and analyzed molecular ex vivo assays. J.G.S., M.L., H.B.S., Ge.B., F.J.T., A.Ö.Y., P.K., M.M.H and M.K. contributed resources. All authors read and revised the manuscript.

## Acknowledgments

We gratefully acknowledge the provision of human biomaterial and clinical data from the CPC-M bioArchive and its partners at the Asklepios Biobank Gauting, the LMU Hospital and the Ludwig-Maximillians-Universität München. We thank the patients and their families for their support. We gratefully acknowledge technical support from Katrin Federl. We are thankful to Roxana M Wasnick for helpful discussions. We further thank A. Feuchtinger and the Core Facility Pathology and Tissue Analytics at Helmholtz Munich for their microscopy and imaging support and Helmholtz AI for support with the image alignment tool. ML, AÖY, HBS acknowledge support from the German Center for Lung Research (DZL) and the Helmholtz Association of German Research Centers. ML acknowledges support from the Deutsche Forschungsgemeinschaft (DFG, German Research Foundation; 512453064) and the von Behring Röntgen Foundation (71_0011). JGS has received funding from the European Respiratory Society and the European Union’s H2020 research and innovation programme under the Marie Sklodowska-Curie RESPIRE4 (grant agreement no. 847462 to JGS and HBS). The cytokeratin-8 antibody (TROMA-I) developed by Brulet, P. / Kemler, R. (Institut Pasteur) was obtained from the Developmental Studies Hybridoma Bank, created by the NICHD of the NIH and maintained at The University of Iowa, Department of Biology, Iowa City, IA 52242.

## Data Supplement

**Supplementary Figure 1.**
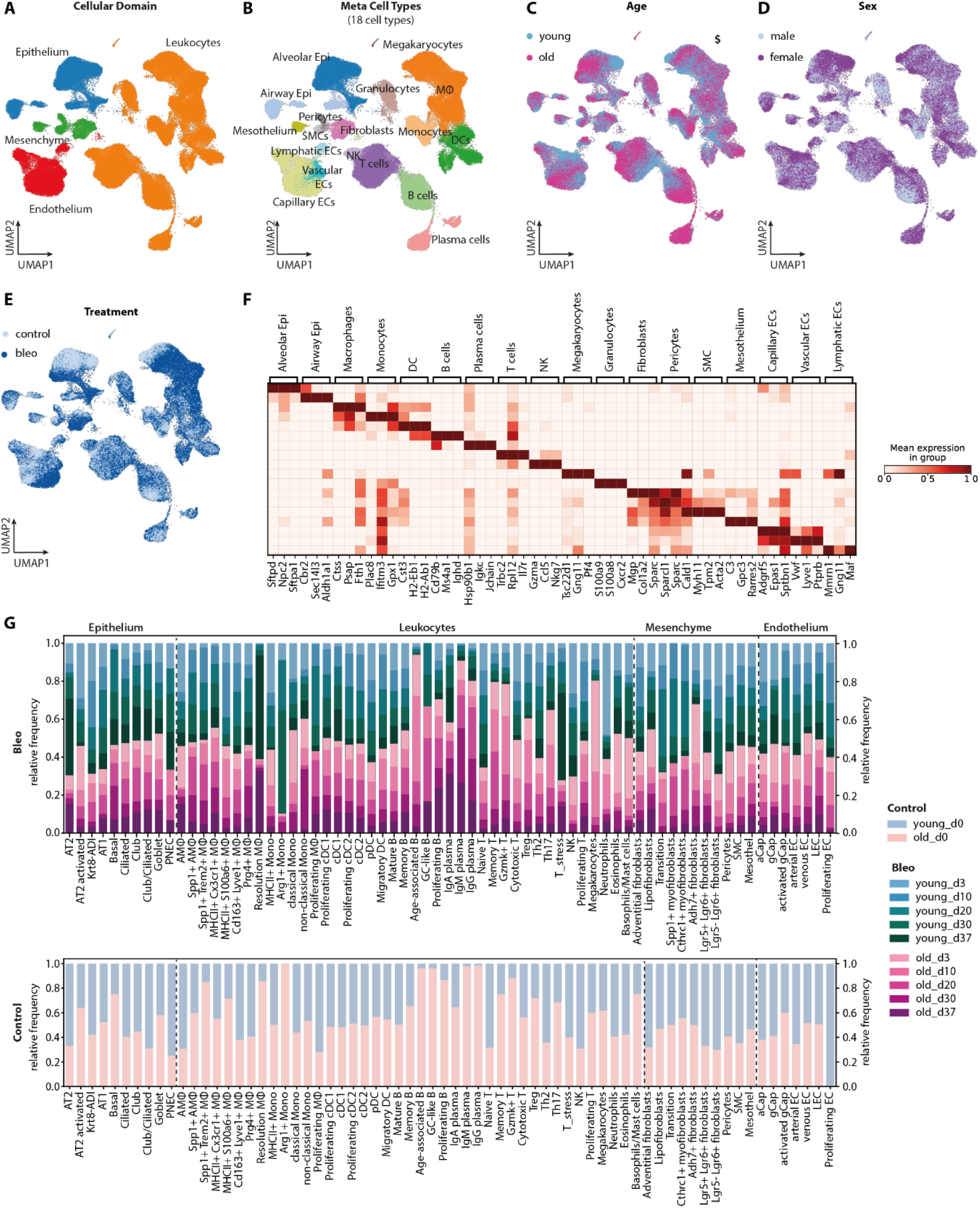
A longitudinal scRNA-seq atlas reveals age-dependent changes in cellular regeneration dynamics following bleomycin-induced lung injury. (A-E) UMAP embedding of the complete single-cell dataset (160,477 cells) colored by cellular domain (A), meta cell type label (B), age (C), sex (D), and treatment group (E). (F) Marker gene signatures of meta cell type populations. Scaled average expression per meta cell type is shown. (G) Time-resolved compositional analysis of cell types/states in lungs from young and old mice at day 0 (lower panel) and after bleomycin injury (top panel).

**Supplementary Figure 2.**
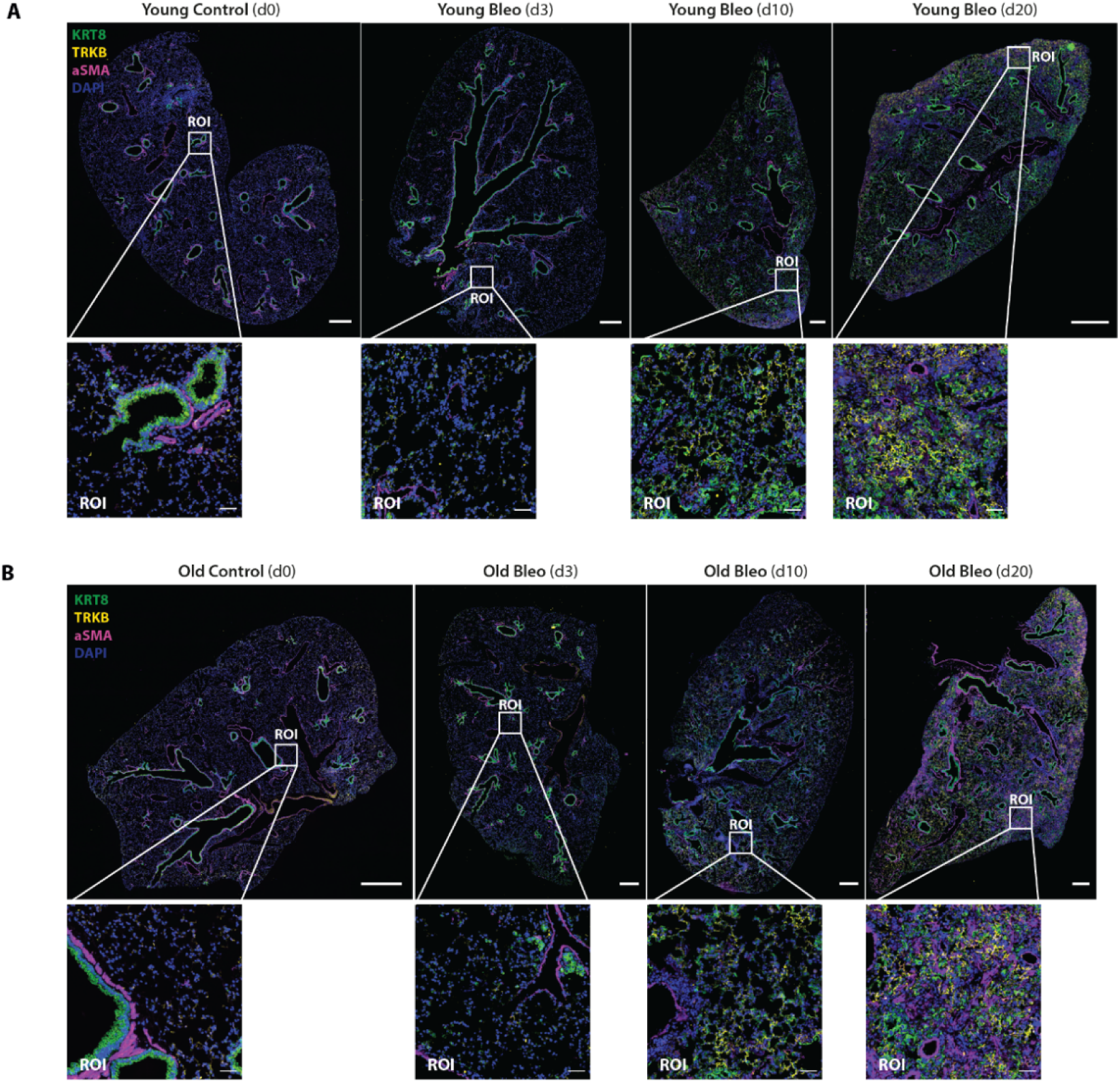
Multiplex immunofluorescence (4i) of epithelial, stromal and endothelial regenerative cell states in young vs aged lungs. (A,B) Representative 4i images of lung tissues from young (A) and old mice (B) at 0, 3, 10 and 20 dpi showing the time-dependent emergence of Krt8-ADI cells, TRKB⁺ activated gCap cells, and αSMA⁺ myofibroblasts following lung injury (scale bar for overview image = 1 mm; ROI scale bars = 50 µm). n = 3 biological replicates.

**Supplementary Figure 3.**
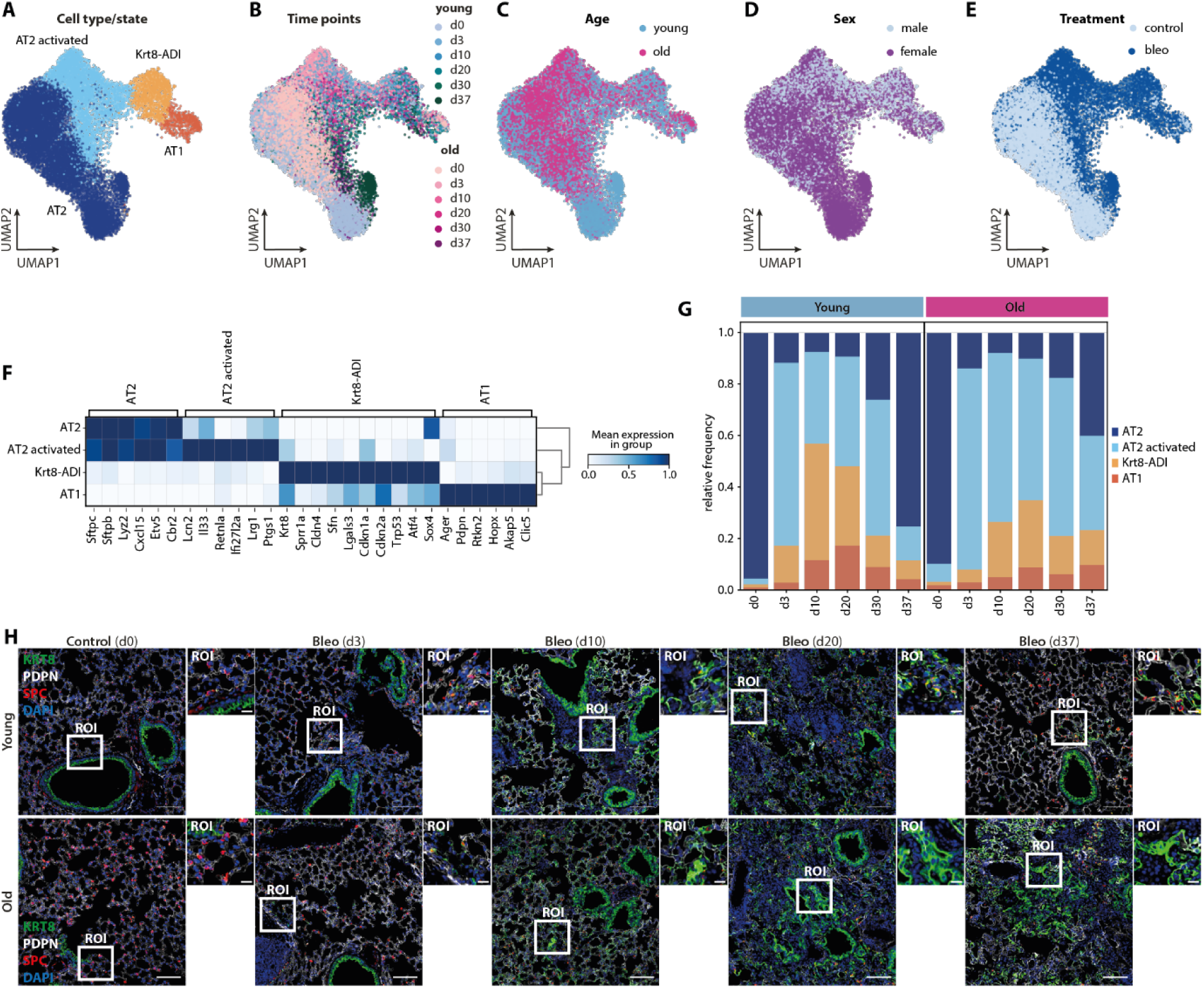
Age-dependent changes in alveolar cell dynamics following bleomycin-induced lung injury. (A-E) UMAP embedding of the alveolar single-cell dataset (19,746 cells) colored by cell type/state annotation (A), time points (B), age (C), sex (D), and treatment group (E). (F) Marker gene signatures of alveolar cell type populations. Scaled average expression per cell type is shown. (G) Time-resolved compositional analysis of alveolar cell types/states in lungs from young and old mice. (H) Representative immunofluorescent images of lung tissues from young (top panel) and old mice (lower panel) at 0, 3, 10, 20 and 37 dpi stained for KRT8 (green), PDPN (white) and pro-SPC (red). Scale bar for overview image = 100 µm; ROI = 20 µm. n = 3 biological replicates.

**Supplementary Figure 4.**
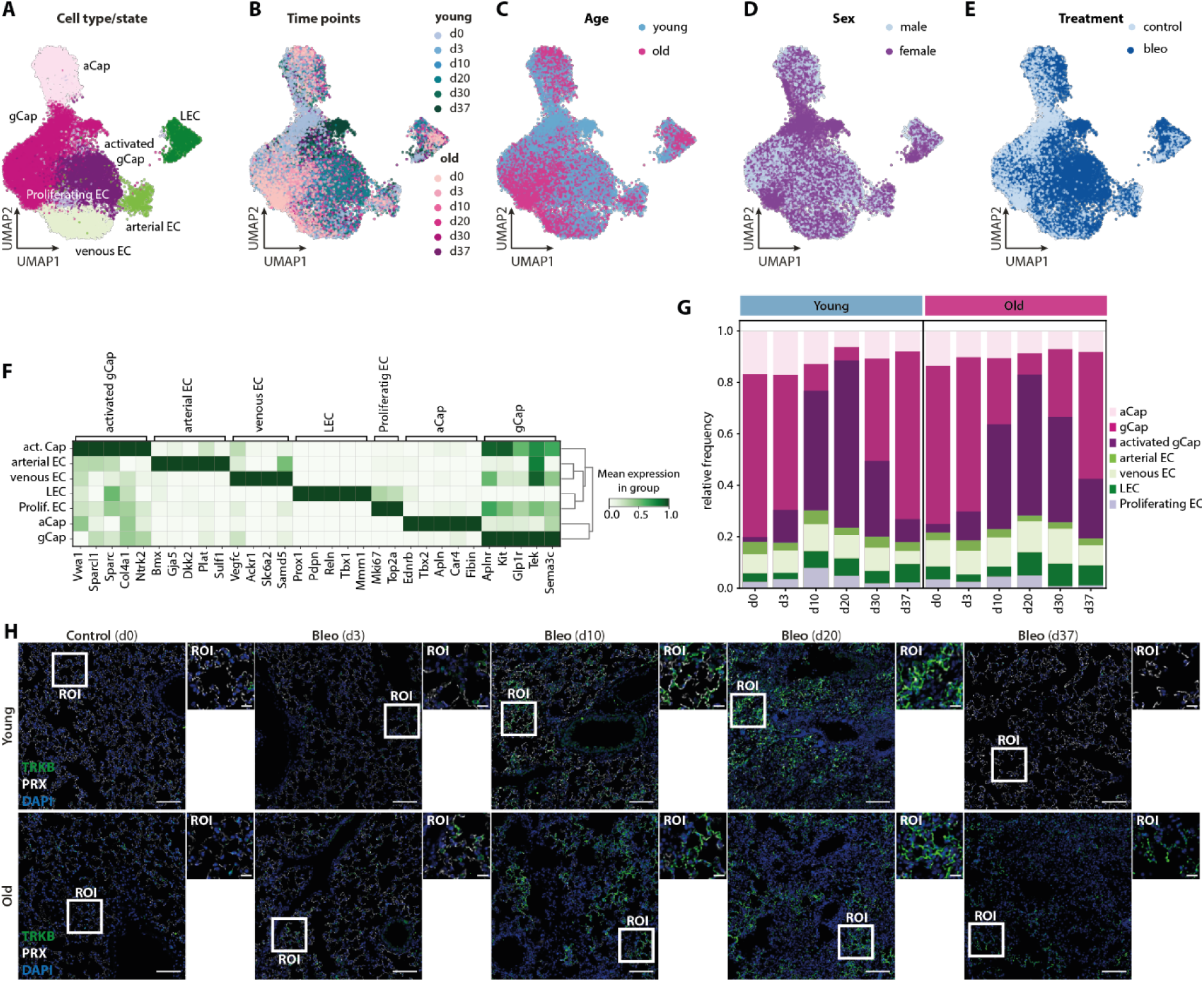
Age-dependent changes in endothelial cell dynamics following bleomycin-induced lung injury. (A-E) UMAP embedding of the endothelial single-cell dataset (24,264 cells) colored by cell type/state annotation (A), time points (B), age (C), sex (D), and treatment group (E). (F) Marker gene signatures of alveolar cell type populations. Scaled average expression per cell type is shown. (G) Time-resolved compositional analysis of endothelial cell types/states in lungs from young and old mice. (H) Representative immunofluorescent images of lung tissues from young (top panel) and old mice (lower panel) at 0, 3, 10, 20 and 37 dpi stained for TRKB (green) and PRX (white). Scale bar for overview image = 100 µm; ROI = 20 µm. n = 3 biological replicates.

**Supplementary Figure 5.**
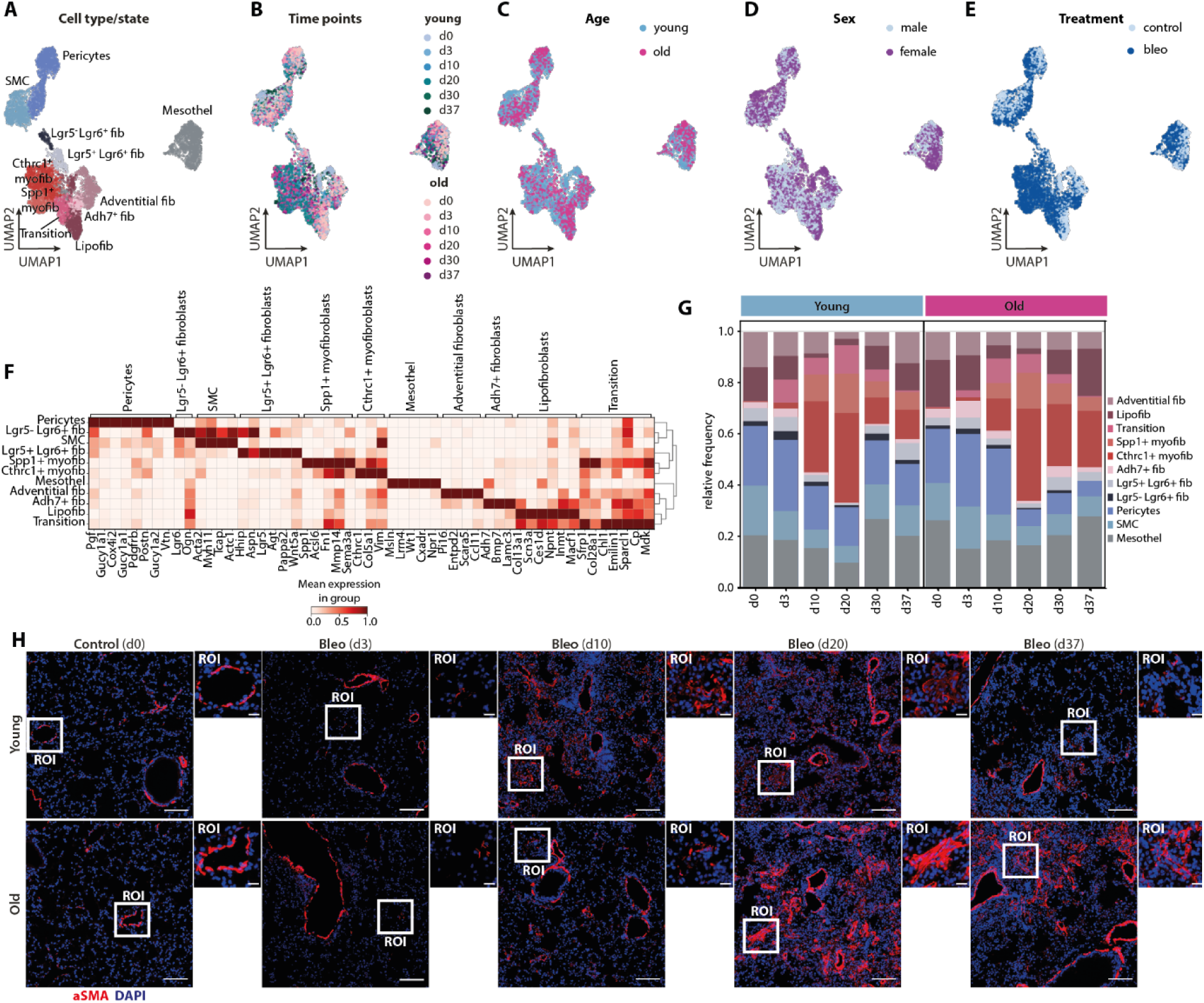
Age-dependent changes in mesenchymal cell dynamics following bleomycin-induced lung injury. (A-E) UMAP embedding of the endothelial single-cell dataset (5,886 cells) colored by cell type/state annotation (A), time points (B), age (C), sex (D), and treatment group (E). (F) Marker gene signatures of mesenchymal cell type populations. Scaled average expression per cell type is shown. (G) Time-resolved compositional analysis of mesenchymal cell types/states in lungs from young and old mice. (H) Representative immunofluorescent images of lung tissues from young (top panel) and old mice (lower panel) at 0, 3, 10, 20 and 37 dpi stained for aSMA (red). Scale bar for overview image = 100 µm; ROI = 20 µm. n = 3 biological replicates.

**Supplementary Figure 6.**
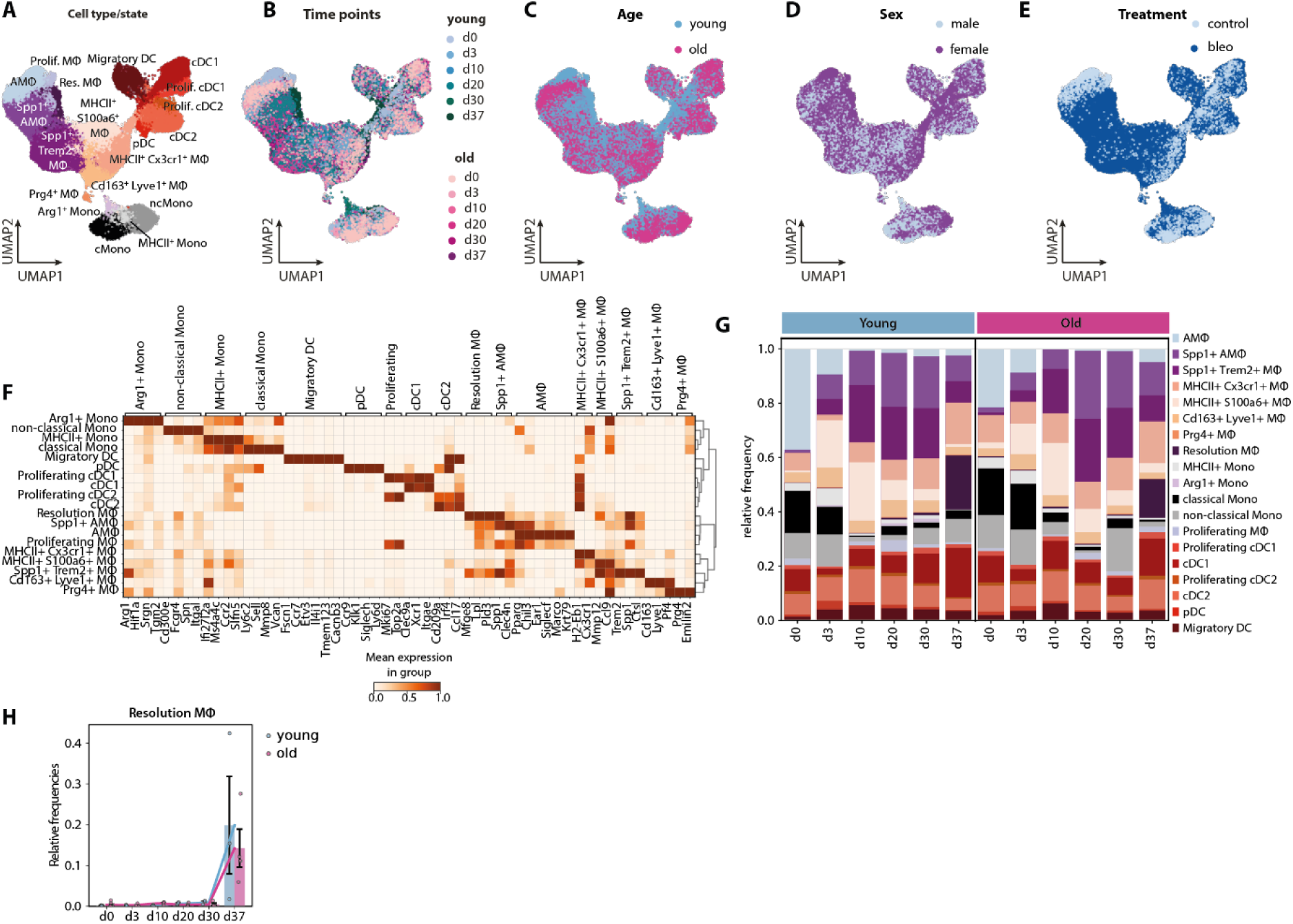
Age-dependent changes in myeloid cell dynamics following bleomycin-induced lung injury. (A-E) UMAP embedding of the myeloid single-cell dataset (52,052 cells) colored by cell type/state annotation (A), time points (B), age (C), sex (D), and treatment group (E). (F) Marker gene signatures of myeloid cell type populations. Scaled average expression per cell type is shown. (G) Time-resolved compositional analysis of myeloid cell types/states in lungs from young and old mice. (H) Time-resolved changes in the relative abundance of resolution macrophages in young vs aged mice. Mean trajectories and individual sample bar plots with standard error of the mean (SEM) are shown.

**Supplementary Figure 7.**
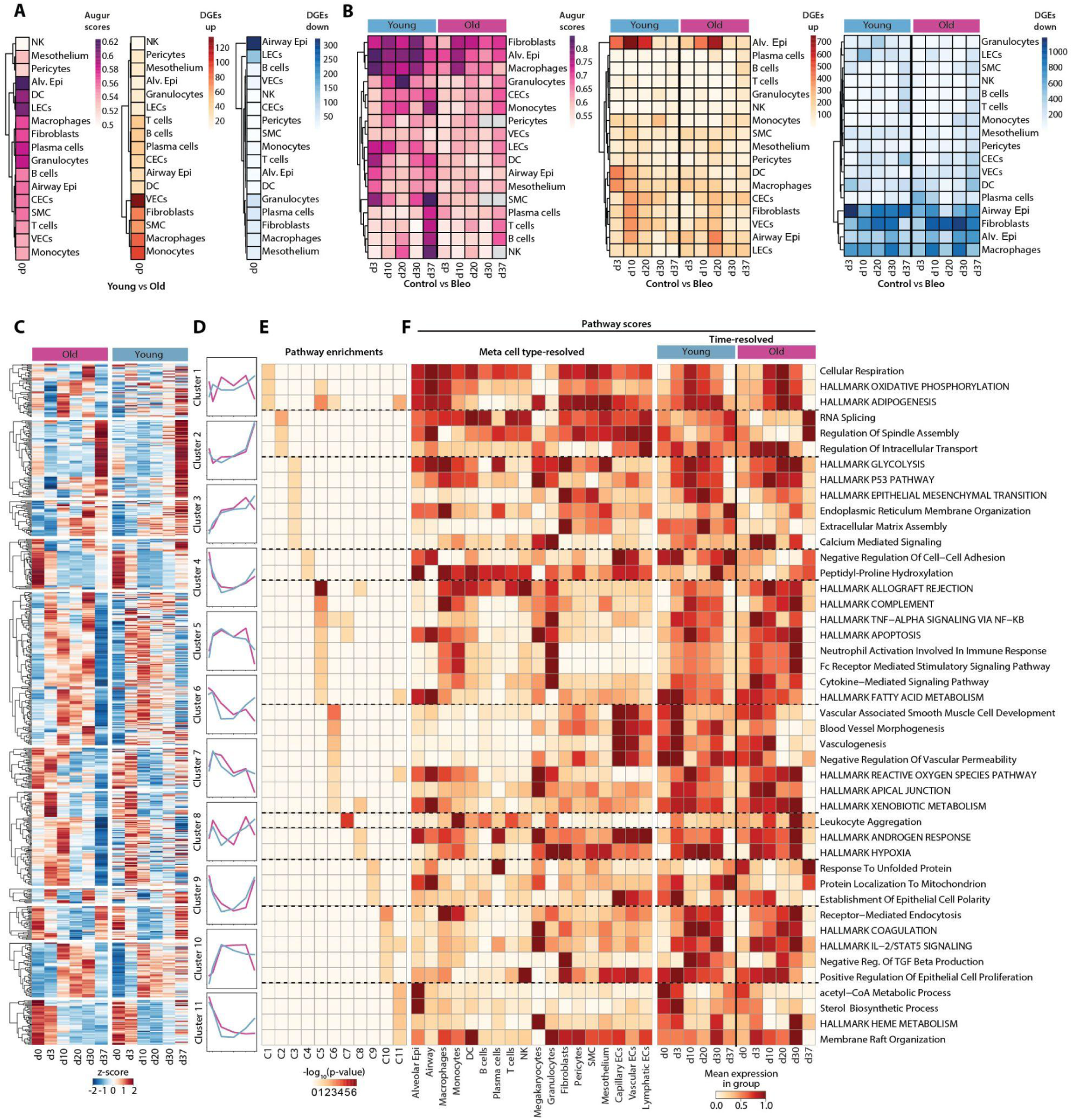
Aging alters molecular programs underlying injury resolution. (A) Heatmap of Augur scores and the number of up- and downregulated genes in young versus aged mice at day 0. (B) Time- and age-resolved heatmap of Augur scores and the number of up- and downregulated genes in control versus bleomycin-treated mice. (C, D) Unsupervised clustering of temporal gene expression profiles based on the 658 downregulated DEGs between young and aged mice at 37 dpi, analyzed at the pseudobulk level. (C) Heatmap showing gene expression profiles across the 11 identified clusters. (D) Average gene expression trajectories of the 11 clusters over time. (C) GO enrichment analysis of cluster-specific gene profiles. (D) Time-resolved gene signature scoring of the identified pathways at the meta cell type level. The scaled average expression is shown.

**Supplementary Figure 8.**
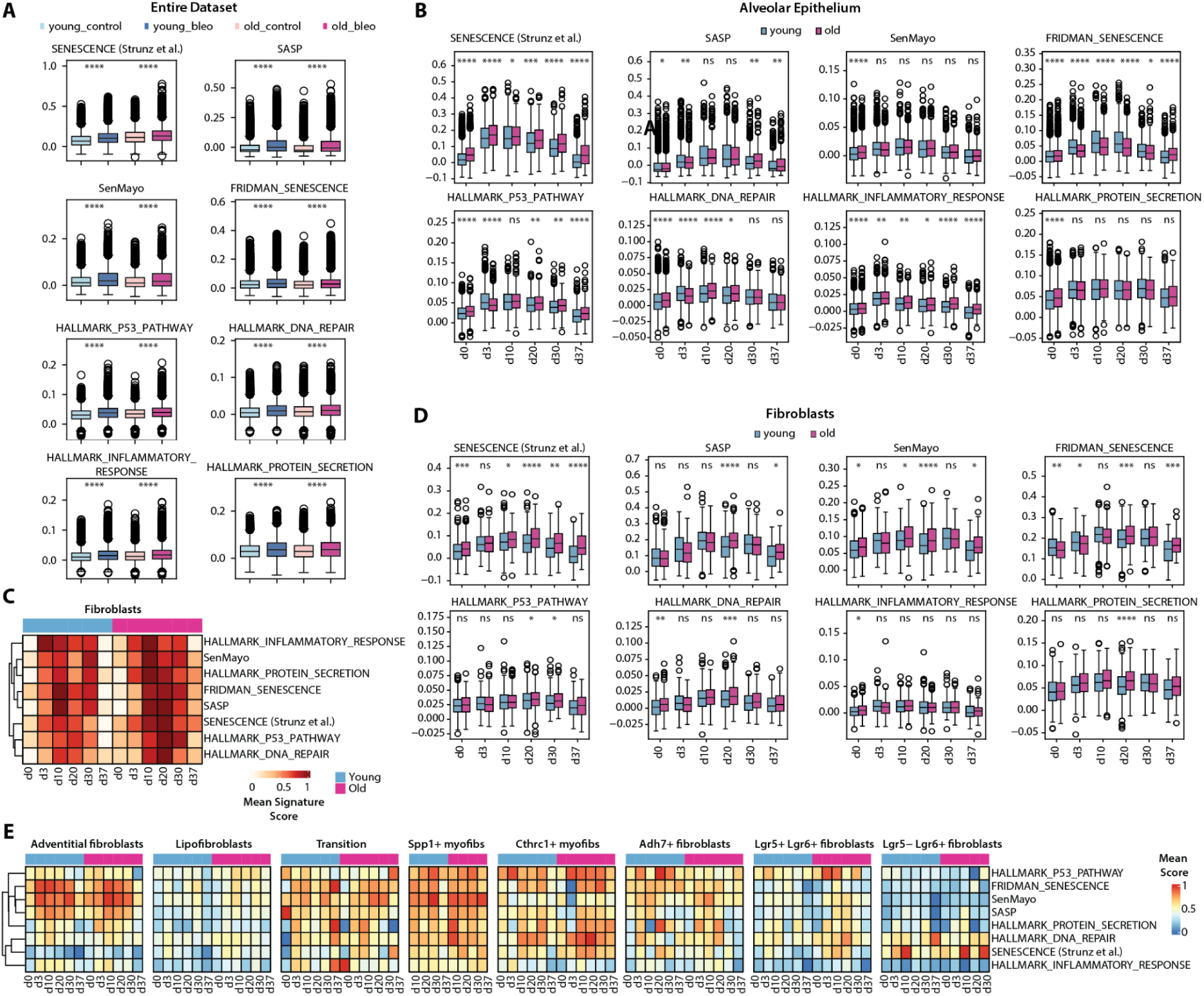
Aging is associated with persistent alveolar and fibroblast senescence following lung injury. (A) Boxplots showing senescence-associated gene signature scores across the entire dataset, stratified by treatment and age. (B) Time- and age-resolved boxplots of senescence-associated gene signature scores within the alveolar epithelial cell compartment. (C–E) Senescence signature analysis within the fibroblast and alveolar epithelial cell compartments. (C) Heatmap showing time- and age-resolved scoring of senescence-associated gene signatures in fibroblasts. (D) Corresponding boxplots for fibroblasts. (E) Heatmap showing signature scores for individual fibroblast cell states over time and by age. Statistics: Mann-Whitney U test.

**Supplementary Figure 9.**
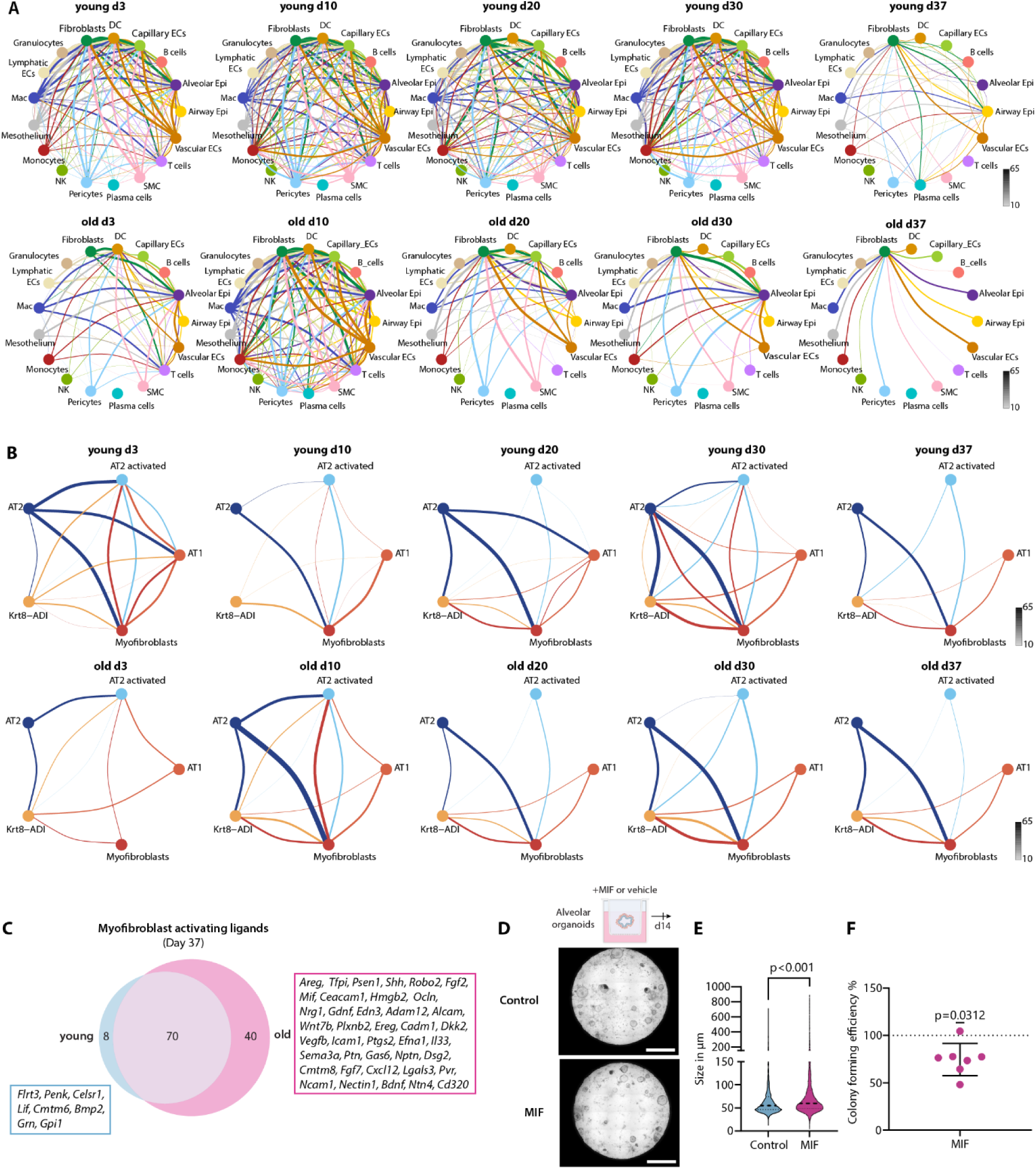
Aging is linked to persisting Krt8-ADI-Myofibroblast crosstalk. (A, B) NicheNet-based analysis of injury-induced cell–cell communication at day 37 post-injury comparing young and aged mice, shown at the meta cell type level (A) and specifically between alveolar epithelial cell states and myofibroblasts (B). (C) Venn diagram showing overlap of predicted myofibroblast-activating ligands identified in young versus aged mice. (D) Representative brightfield images of lung organoid cultures treated with vehicle or recombinant MIF. Scale bars = 50 µm. (E, F) Quantification of organoid sizes (µm) (E) and colony-forming efficiency (%) (F) in cultures treated with MIF versus vehicle. Statistical analysis: paired Wilcoxon test; *n* = 7 biological replicates.

**Supplementary Figure 10.**
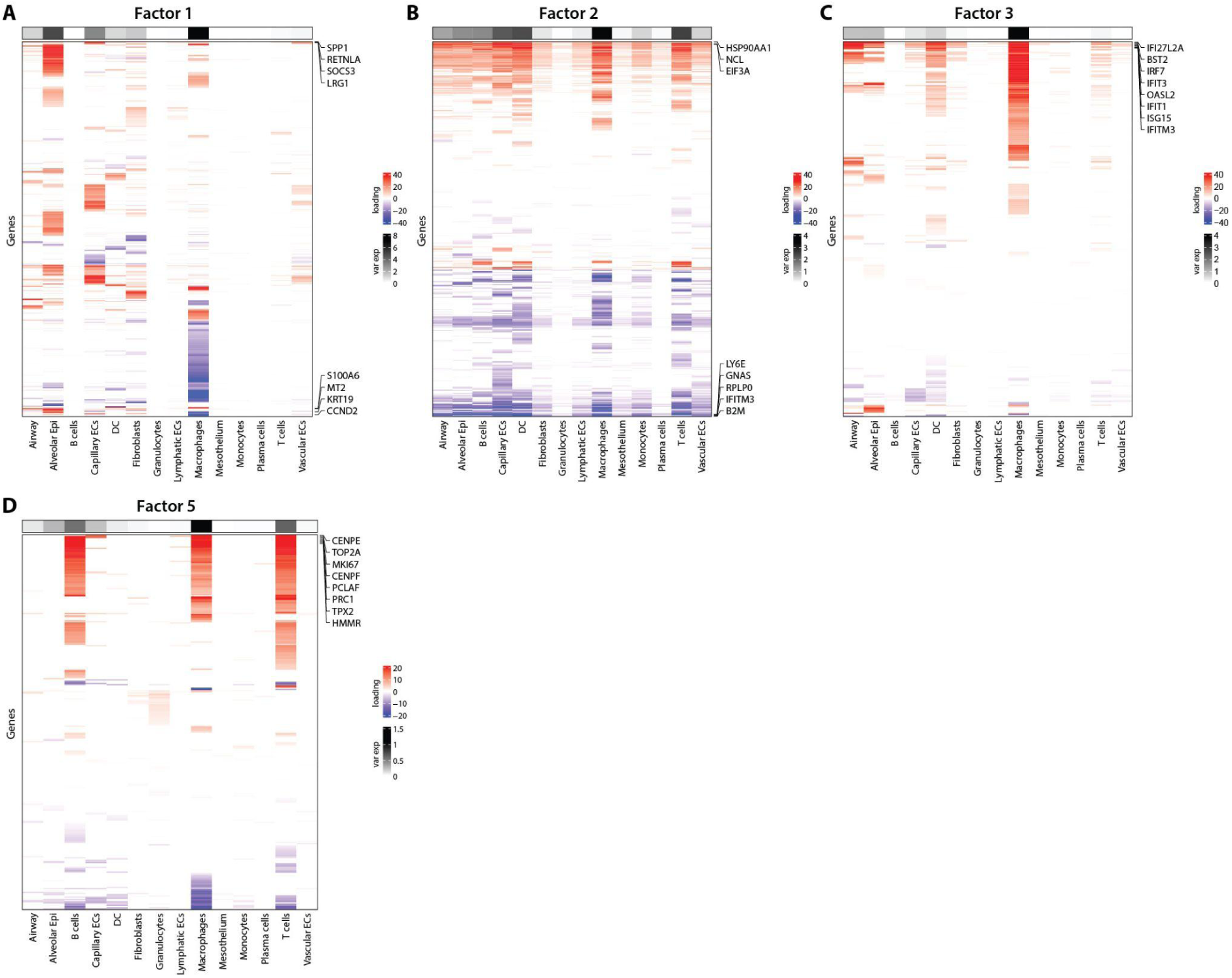
scITD-based multicellular gene program analysis. (A–D) Loading matrices for scITD factors 1 (A), 2 (B), 3 (C), and 5 (D), highlighting key gene contributors to each multicellular transcriptional program. (E-G) Cell type contribution to the scITD factors 2 (E), 3 (F) and 5 (G)

**Supplementary Figure 11.**
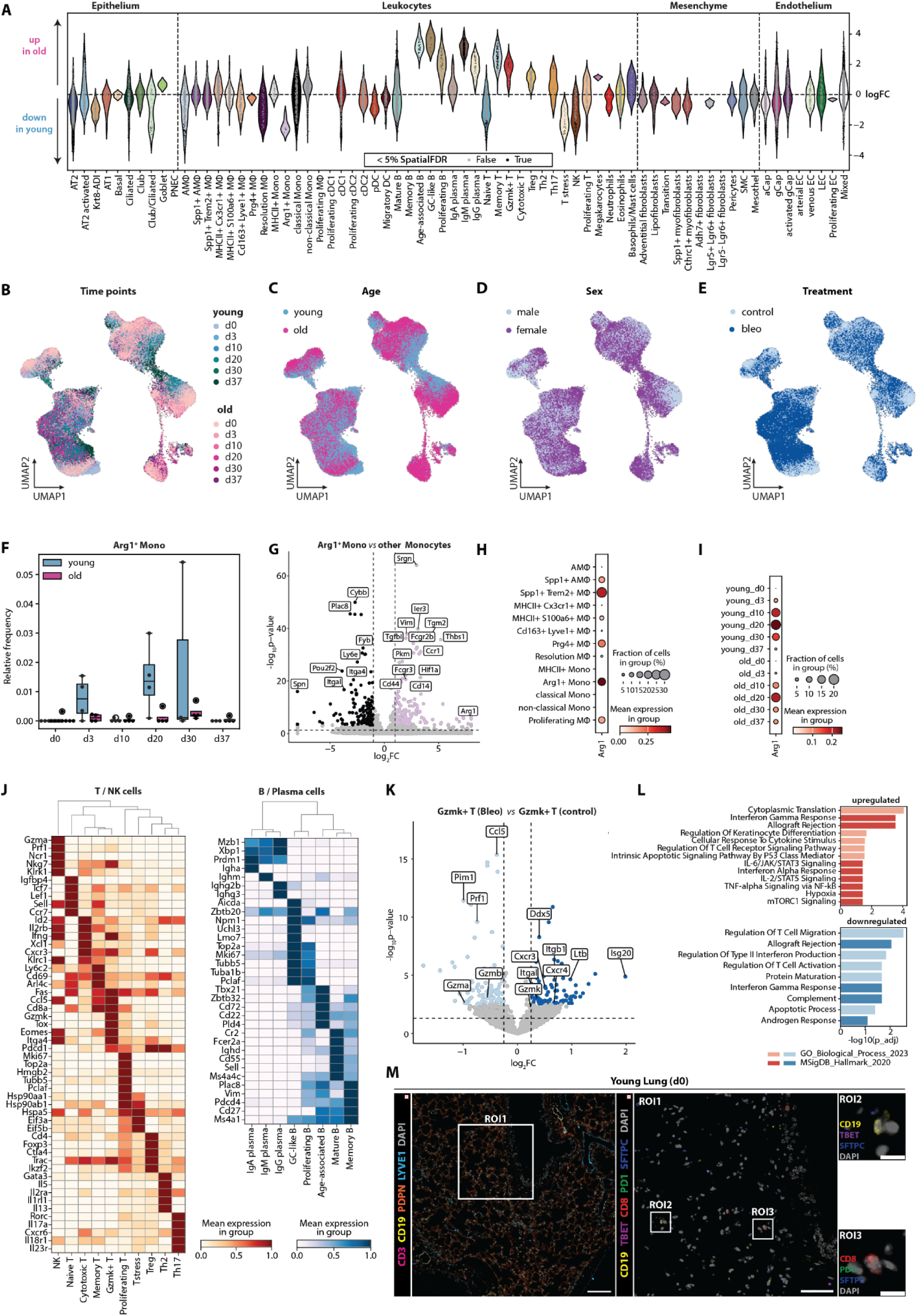
Lung aging is associated with immune compartment remodeling and the emergence of age-specific lymphocyte populations. (A) Milo analysis of differential cellular neighborhoods in the full dataset comparing young and aged lungs. Beeswarm plot displays differentially abundant neighborhoods mapped to cell type annotations, showing log-fold changes in young versus aged mice. (B–E) UMAP representations of the macrophage/monocyte and T/B lymphocyte compartments colored by time point (B), age (C), sex (D), and treatment group (E). (F) Time-resolved changes in the relative abundance of Arg1⁺ monocytes in young and aged mice. (G) Volcano plot showing differentially expressed genes (DEGs) between Arg1⁺ monocytes and other monocyte populations. (H, I) Dot plots showing mean expression of Arg1 in young versus aged mice, resolved by cell type (H) or over time (days 0–37, I). (J) Marker gene signatures of T/NK and B/plasma cell populations. Scaled average expression per cell type is shown. (K) Volcano plot displaying DEGs between bleomycin-treated and untreated Gzmk⁺ T cells in aged mice. (L) Gene Ontology (GO) enrichment analysis of upregulated (top panel) and downregulated (bottom panel) pathways in bleomycin-treated versus untreated Gzmk⁺ T cells in aged mice. (M) Representative 4i image of lung tissue from young mice showing no accumulation of CD19⁺/TBET⁺ age-associated B cells or CD8⁺/PD1⁺ Gzmk⁺ T cells, and absence of tertiary lymphoid-like structures (scale bar for overview image = 100 µm; ROI1, scale bars = 50 µm). ROI2: zoom-in of typical CD19⁺ B cells located in the alveolar septum. ROI3: zoom-in of CD8⁺ T cells located in the alveolar septa. Scale bars = 10 µm. n = 3 biological replicates.

**Supplementary Figure 12.**
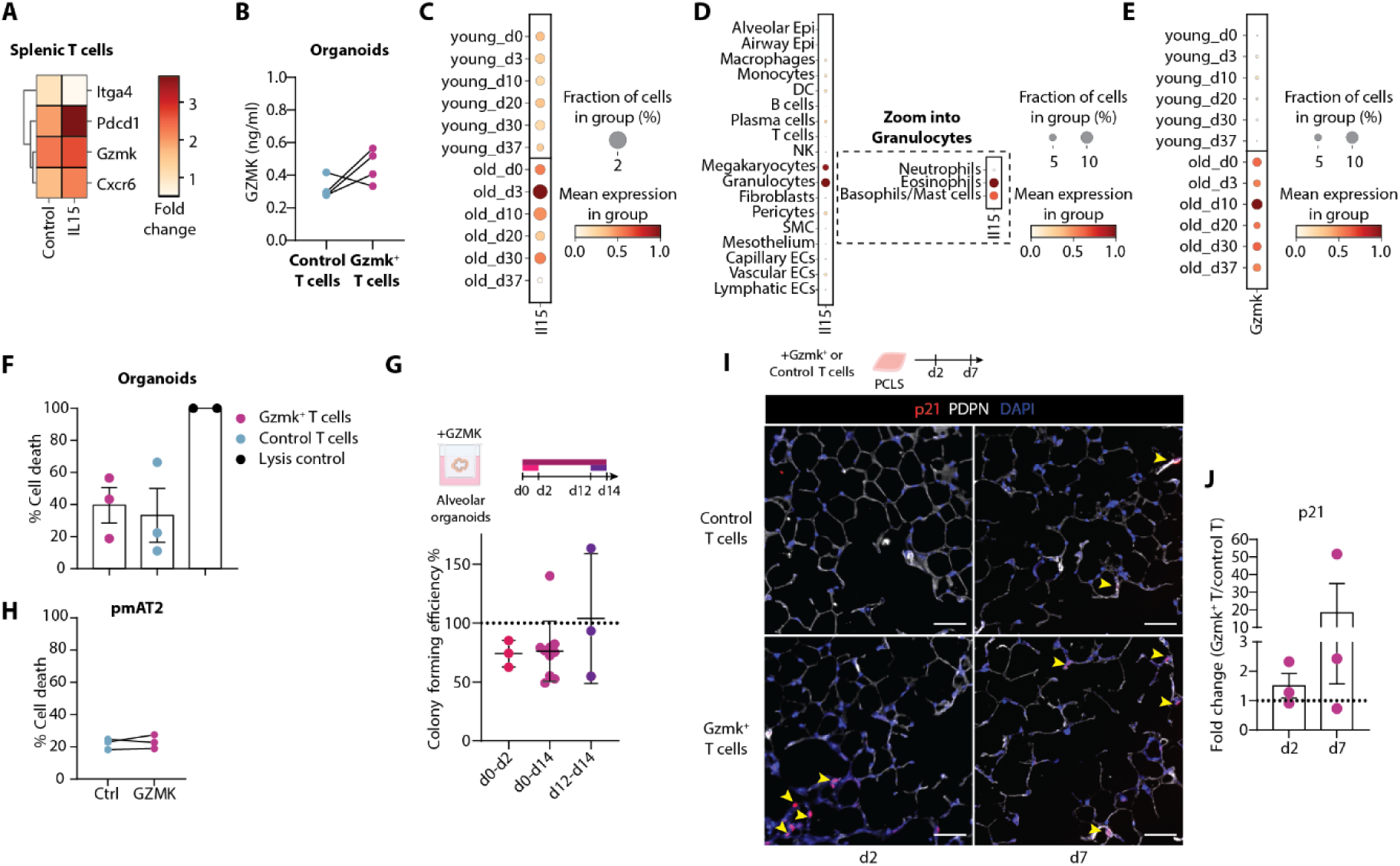
Gzmk⁺ T Cells Impair Alveolar Regeneration via Induction of Epithelial Senescence. (A) Relative fold change in mRNA expression of Gzmk⁺ T cell markers (*Itga4, Cxcr6, Pdcd1, Gzmk*) in IL-15-stimulated splenic T cells from young mice versus controls (n=4 biological replicates). (B) GZMK protein levels (ng/mL) in organoid culture assays on day 14 following co-culture with unstimulated or IL15 stimulated splenic CD8^+^ T cells. (C, D) Dot plots showing mean expression of *Il15* in young versus aged mice, resolved by time (day 0–37, C) or by meta cell type (D). (E) Dot plot of mean Gzmk expression in the lymphoid compartment of young versus aged mice, resolved by time (day 0–37). (F) LDH cytotoxicity assay of alveolar organoid cultures co-cultured with Gzmk⁺ or control T cells (n=3 biological replicates). (G) Colony forming efficiency (%) of alveolar organoids following time-dependent stimulation with recombinant GZMK (n=3-7 biological replicates). (H) LDH cytotoxicity assay of primary murine AT2 cells (pmAT2) stimulated with GZMK or vehicle (n=3 biological replicates). (I) Representative immunofluorescence images showing expression of Cdkn1a/p21 (red) and Podoplanin (PDPN, white) in PCLS co-cultured with Gzmk⁺ or control T cells. Scale bars = 50 µm; n = 3 biological replicates. (J) Quantification of immunofluorescence stainings shown in (I).

